# Structural basis of peptidoglycan endopeptidase regulation

**DOI:** 10.1101/843615

**Authors:** Jung-Ho Shin, Alan G. Sulpizio, Aaron Kelley, Laura Alvarez, Shannon G. Murphy, Felipe Cava, Yuxin Mao, Mark A. Saper, Tobias Dörr

## Abstract

Most bacteria surround themselves with a cell wall, a strong meshwork consisting primarily of the polymerized aminosugar peptidoglycan (PG). PG is essential for structural maintenance of bacterial cells, and thus for viability. PG is also constantly synthesized and turned over, the latter process is mediated by PG cleavage enzymes, for example the endopeptidases (EPs). EPs themselves are essential for growth, but also promote lethal cell wall degradation after exposure to antibiotics that inhibit PG synthases (*e.g.*, β-lactams). Thus, EPs are attractive targets for novel antibiotics and their adjuvants. However, we have a poor understanding of how these enzymes are regulated *in vivo*, depriving us of novel pathways for the development of such antibiotics. Here, we have solved crystal structures of the LysM/M23 family peptidase ShyA, the primary EP of the cholera pathogen *Vibrio cholerae.* Our data suggest that ShyA assumes two drastically different conformations; a more open form that allows for substrate binding, and a closed form, which we predicted to be catalytically inactive. Mutations expected to promote the open conformation caused enhanced activity *in vitro* and *in vivo*, and these results were recapitulated in EPs from the divergent pathogens *Neisseria gonorrheae* and *Escherichia coli*. Our results suggest that LysM/M23 EPs are regulated via release of the inhibitory Domain1 from the M23 active site, likely through conformational re-arrangement *in vivo*.

**Significance:** Bacteria digest their cell wall following exposure to antibiotics like penicillin. The endopeptidases (EPs) are among the proteins that catalyze cell wall digestion processes after antibiotic exposure, but we do not understand how these enzymes are regulated during normal growth. Herein, we present the structure of the major EP from the diarrheal pathogen *Vibrio cholerae.* Surprisingly, we find that EPs from this and other pathogens appear to be produced as a largely inactive precursor that undergoes a conformational shift exposing the active site to engage in cell wall digestion. These results enhance our understanding of how EPs are regulated and could open the door for the development of novel antibiotics that overactivate cell wall digestion processes.

## Introduction

The bacterial cell wall is essential for growth, shape maintenance, and survival of most free-living bacteria. Due to its vital role in bacterial structural integrity, the cell wall is a time-tested target for our most effective antibiotics and continues to be at the center of drug discovery efforts (1). The major constituent of the bacterial cell wall is peptidoglycan (PG), a complex macromolecule composed of polysaccharide strands that are crosslinked via short oligopeptide sidestems to form a mesh-like structure. PG is assembled outside of the cytoplasmic membrane from a lipid-linked disaccharide-pentapeptide precursor called lipid II. During cell wall assembly, lipid II is polymerized via glycosyltransferase activities of class A penicillin-binding proteins (aPBPs) and **s**hape, **e**longation, **d**ivision and **s**porulation (SEDS) proteins RodA and FtsW. The resulting PG strands are then crosslinked with one another via a transpeptidation reaction, catalyzed by the bifunctional aPBPs and by class B PBPs (bPBPs) that associate with SEDS proteins (2–7).

Due to its importance to maintain structural integrity in the face of changing environmental conditions that impose variable osmotic pressures, PG (in Gram-negative bacteria in conjunction with the outer membrane (8)) must form a tight and robust cage around the bacterial cell. While maintaining the cell wall’s vital structural role, however, the cell must also be able to expand (beyond the limits of what stretching of PG could accomplish (9, 10)), divide, and insert macromolecular protein complexes into its cell envelope. Thus, cell wall synthesis alone is insufficient; PG must also be constantly modified, degraded and resynthesized. These turnover processes are mediated by a group of enzymes often collectively referred to as “autolysins”. Autolysins cleave a variety of bonds in the PG meshwork and in doing so fulfill important physiological functions such as shape generation (endopeptidases, EPs (11, 12)) daughter cell separation (amidases and lytic transglycosylases, LTGs (13–19)), insertion of trans-membrane protein complexes (LTGs, EPs (20–22)), and cell elongation (EPs (23–26)).

Because they can cleave bonds within the PG meshwork, autolysins are potentially as dangerous as they are important, since unchecked cell wall degradation can in principle result in cell lysis and death. Amidases (which cut between polysaccharide backbone and peptide sidestem), LTGs (which cleave the PG polysaccharide backbone) and EPs (which cleave the bond between peptide sidestems), for example, degrade PG following exposure to β-lactam antibiotics (15, 27–30). This results in cell lysis or the formation of non-dividing, cell wall-deficient spheroplasts (27, 31, 32). Endopeptidases play a particularly dominant role during antibiotic-induced spheroplast formation, at least in the cholera pathogen *Vibrio cholerae* (27). Autolysin genes are also often highly abundant. *V. cholerae*, for example, encodes nine predicted endopeptidases, of which only ShyA and ShyC are collectively essential for cell elongation (23). Similarly, the model organism *E. coli* encodes at least 8 EPs (24, 33, 34). Due to this inherently perilous potential, endopeptidases must be tightly regulated under normal growth conditions to ensure proper PG turnover without compromising structural integrity.

We currently have an incomplete understanding of how endopeptidases are regulated in bacteria. The Gram-positive bacterium *Bacillus subtilis* (and likely other Gram-positive species as well) regulates EP activity via interactions with FtsEX and the PG synthesis elongation machinery (35–37) and additionally encodes a post-translational negative regulator, IseA (YoeB) (38), which presumably serves to tone down EP activity during cell wall stress conditions (38, 39). Much less is known about EP regulation in Gram-negative bacteria, where EPs appear to be kept in check as part of multiprotein complexes (40, 41), at the transcriptional level (42, 43) or via proteolytic degradation (44, 45). Proteolytic turnover appears to be the major mode of regulation of growth-promoting EPs (44, 45); however, phenotypes associated with the accumulation of EPs have surprisingly mild phenotypes under normal growth conditions, suggesting an additional, unexplored layer of regulation.

Here, we present two crystal structures of the major EP ShyA of *V. cholerae*. The ShyA structure yielded two drastically different conformations, of which only one appears to be active. We thus propose that EPs are produced in a predominantly inactive form and regulated by stabilizing their active conformation *in vivo.* Importantly, the domain organization and mechanism of regulation via conformational switching appears to be conserved among diverse Gram-negative pathogens. These results expose a new mechanism of endopeptidase regulation in Gram-negative pathogens that could open up new avenues for the development of novel antibiotics that activate EPs to the detriment of bacterial viability.

## Results

### Genetic evidence suggests that LysM/M23 endopeptidases are produced in an inactive form

The ability of EPs to efficiently degrade peptidoglycan poses an exceptional danger to cells. Indeed, purified ShyA almost completely digests purified PG sacculi *in vitro* (23) through its D,D-endopeptidase activity (42) and the EPs play a crucial role in cell wall degradation as a downstream consequence of exposure to β-lactam antibiotics (27, 30). Interestingly, we found that overproduction of *V. cholerae’s* two principal LysM/M23 EPs did not cause any obvious adverse growth phenotypes (**Fig. 1A**). Functional expression of both EPs was verified either by Western blot (ShyA only, **Fig. S1A**) or via the ability to complement an EP depletion mutant (ShyA and ShyC, (42)). Thus, EP-mediated PG cleavage activity is either tightly regulated *in vivo* or always outpaced by a highly effective cell wall synthesis machinery. However, we also noticed during depletion experiments of IPTG-inducible ShyA in a background deleted in all other M23 and P60 family endopeptidases (Δ6 endo, i.e. Δ*shyABC*Δ*vc1537*Δ*tagE1,2* P_IPTG_:*shyA*) that viability declined and morphological defects (compared to a control culture that contained inducer) materialized rapidly (within 1 hour) upon subculturing into medium without inducer (**Fig. 1B,C**). Western Blot analysis revealed that at these early time points, ShyA protein levels were still clearly detectable (**Fig. 1D**). Based on these results, we speculated that ShyA might be produced in a predominantly inactive form *in vivo.* We also estimated native ShyA levels using semi-quantitative Western Blot with purified protein of known concentration as a calibration standard. We estimate that under exponential growth conditions in rich medium, ShyA is present at a level of ∼1500 molecules/cell (**Fig. S1B**).

**Figure 1.**
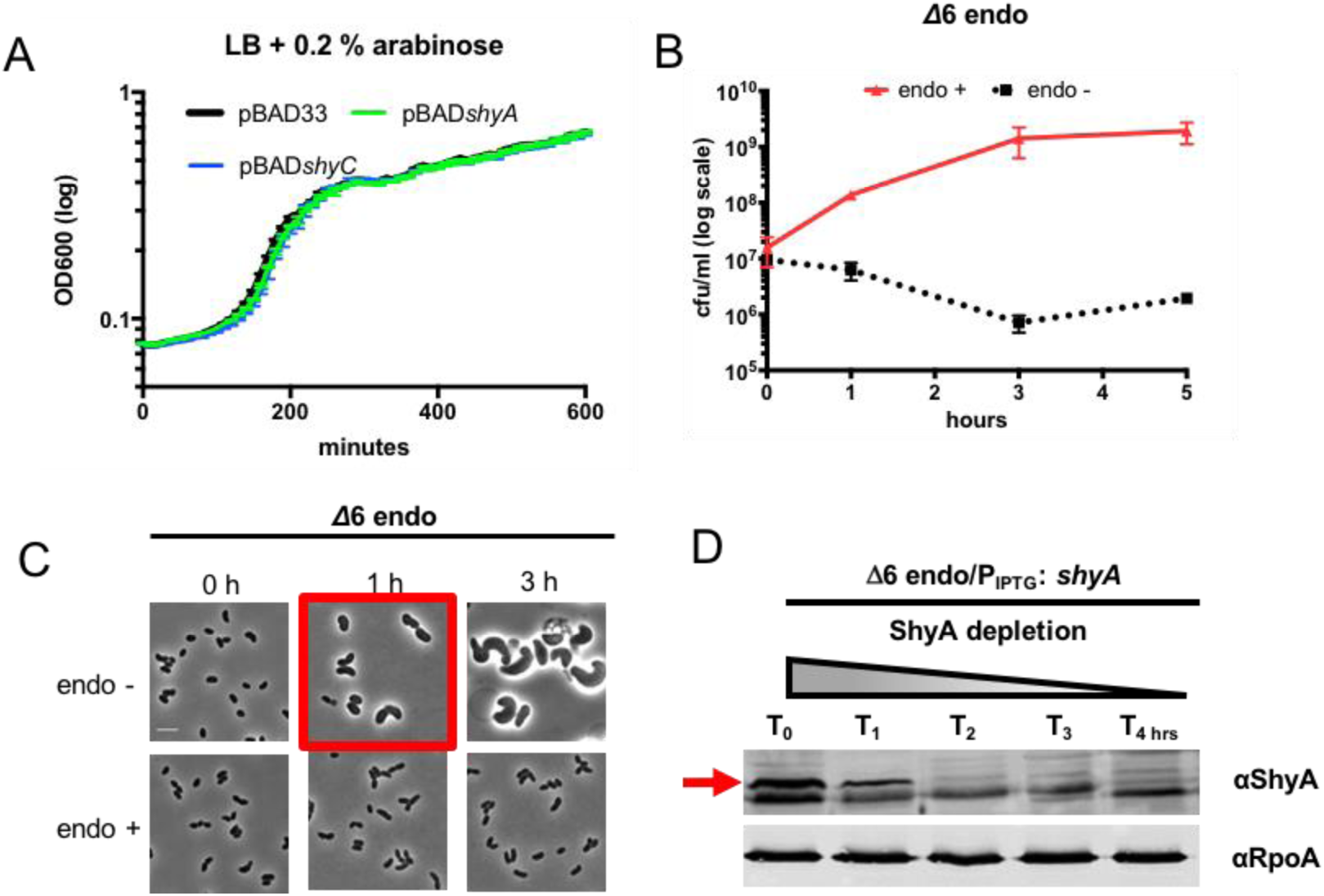
ShyA overexpression and depletion phenotypes. **(A)** EP overexpression does not affect growth behavior. Strains carrying either empty pBAD or EPs under control of an arabinose-inducible promoter were grown in a Bioscreen growth curve analyzer. **(B)** and **(C)** Defects associated with EP depletion materialize early in the depletion timecourse. A Δ6 endo strain (Δ*shyABC* Δ*vc1537* Δ*tagE1/2* P_IPTG_:*shyA*) was grown overnight in the presence of 200 µM IPTG, washed twice and resuspended either in the absence (“endo -“) or presence (“endo +”) of inducer. At designated time points, cells were then either plated on LB agar containing IPTG **(B)**, or imaged on an agarose pad **(C)**. Data in **(B)** are averages of 3 independent experiments, error bars represent standard deviation. **(D)** Western Blot analysis of *in vivo* ShyA levels during early depletion time points. A Δ6 endo/Flag strain (P_IPTG_:*shyA-flag*^sw^) was treated as described under **(B)**. At designated time points, ShyA was visualized via Western Blot using anti-Flag primary antibody. ShyA::Flag is *shyA-flag* expressed from its native locus, +endo is the Δ6 endo/Flag strain grown for 3 h in the presence of IPTG.

### The ShyA crystal structure reveals a putative switch from an inactive to an active conformation

Since our genetic data hinted towards the existence of an inactive form of ShyA *in vivo,* we used a structural approach to probe the mechanistic reason for ShyA putative inactivity. We purified and crystallized two constructs, an N-terminally tagged version (replacing the native signal sequence with a 6xHis-tag) and one tagged at its C-terminus (likewise with a deleted signal sequence). These two constructs serendipitously yielded two different structures, as detailed below.

### Structure of ShyA^CLOSED^

The tertiary structure of ShyA obtained from the monoclinic crystals grown from the N-terminal His-tagged protein was solved by molecular replacement using the previously determined ShyB structure [PDB entry 2GUI (46)] to a resolution of 2.11 Å with good crystallographic statistics (**Table 1**). The solved structure revealed three domains common in other M23 family endopeptidases (46–48) (**Fig. 2A**). Domain 1 contains a LysM PG binding domain (49). Domain 2 appears to serve as a hinge domain between Domain 1 and 3, and further engages in interactions with a C-terminal α-helix. Finally, Domain 3 contains the M23 domain responsible for cleavage of the peptide bond crosslink (D-Ala–diaminopimelic acid [DAP]) in peptidoglycan. Domain 1 and Domain 3 have both hydrophobic and electrostatic interactions bringing the two domains into close proximity. We will henceforth refer to this structure as ShyA^CLOSED^ (**Fig. 2B**).

**Figure 2.**
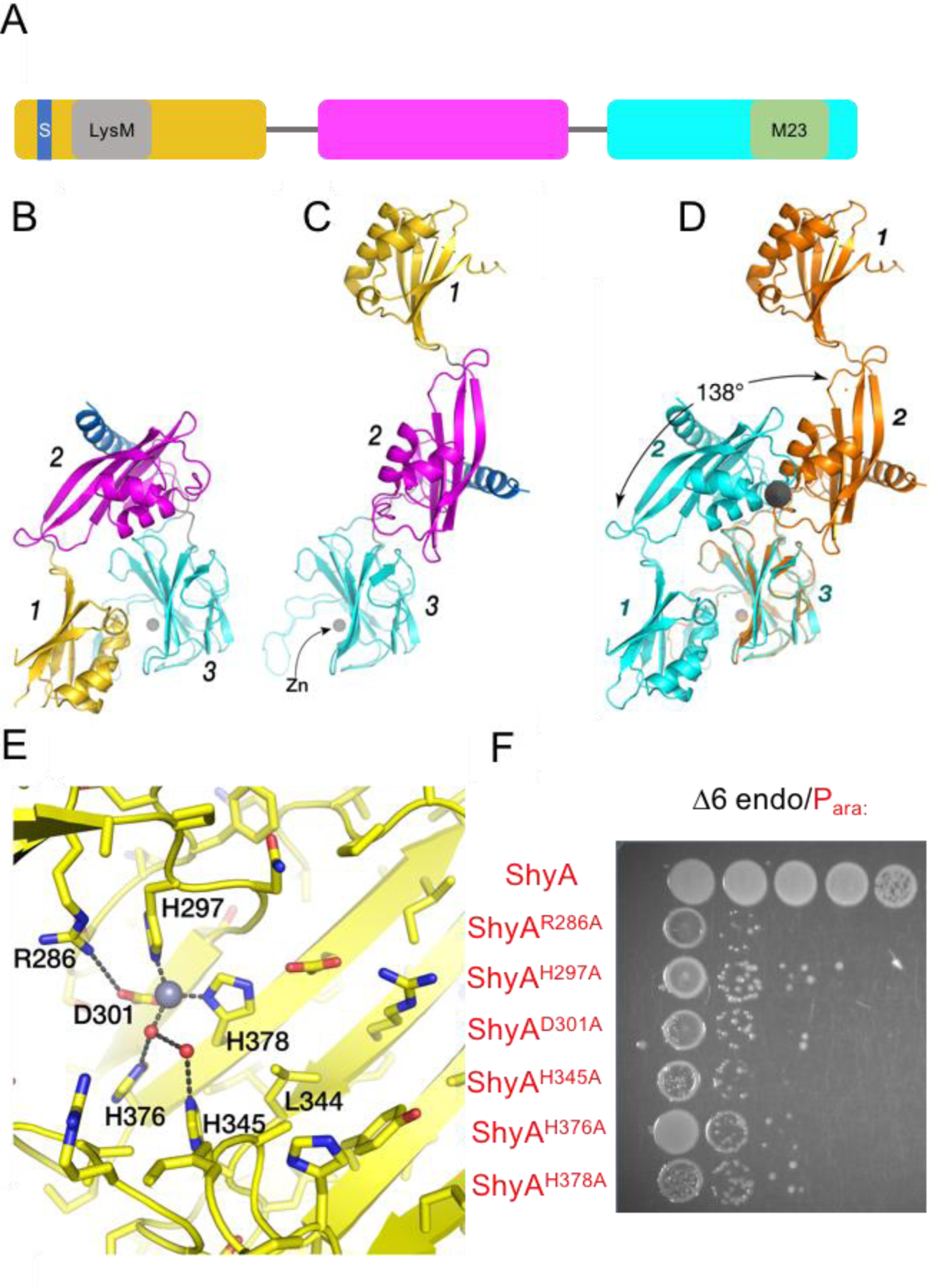
Comparison of the two observed conformations of ShyA. **(A)** Diagram of predicted ShyA domains. The signal sequence (S), LysM domain and M23 catalytic domain are indicated (**B)** Cartoon diagram of ShyA^CLOSED^ (PDB ID 6UE4) determined from crystals grown from protein containing an amino terminal 6xHis-tag. Each domain is numbered and highlighted in a different color. **(C)** Cartoon diagram of ShyA^OPEN^ (PDB ID 6U2A) from crystals grown from ShyA protein containing a carboxyl terminal 6xHis-tag. Domain 3 is oriented exactly like Domain 3 in part **(B)**. **(D)** Domain 3 of each structure are superposed. Closed form in cyan, open form in orange. The grey solid circle is an end-on view of the 138° rotation axis that relates Domain 2 of the two different structures, Note that the C-terminal helix packing against Domain 2 is also related by this same transformation. **(E)** Close-up view of the predicted ShyA active site showing residues putatively required for catalytic function. Red spheres represent water molecules, the grey sphere represents the catalytic zinc atom **(F)** Validation of predicted active site residues in a Δ6 endo background (Δ*shyABC* Δ*vc1537* Δ*tagE1/2* P_IPTG_:*shyA*) where ShyA is under IPTG control. The indicated mutants under control of an arabinose-inducible promoter were plated in the absence of IPTG, but in the presence of arabinose to test for functionality of variants when expressed as the sole EP.

**Table 1.** Data collection and refinement statistics.

|  | ShyA<sup>CLOSED</sup>, PDB ID 6UE4 | ShyA<sup>OPEN</sup>, PDB ID 6U2A |
| --- | --- | --- |
| Wavelength (Å) | 0.9775 | 1.078 |
| Space group | *P* 2<sub>1</sub> | *C* 2 |
| Unit cell (Å, °) | 61.12 87.25 74.97<br>90 100.37 90 | 75.27 82.49 82.52<br>90 102.69 90 |
| Resolution range for data processing (Å) | 49.55–2.08 (2.12–2.08) | 54.9–2.03 (2.28–2.03) |
| Resolution cutoffs from anisotropic analysis (Å):<br>0.78 *a* * – 0.63 *c* *<br>*b* *<br>0.49 *a* * + 0.87 *c* * |  | 2.32<br>3.02<br>1.98 |
| Total reflections measured | 309,587 (15,147) | 121,494 (5,375) |
| Unique reflections | 46,207 (2,295) | 17,391 (870) |
| Multiplicity | 6.7 (6.6) | 7.0 (6.18) |
| Completeness (%)<br>spherical<br>ellipsoidal |  | 54.1 (9.2)<br>89.8 (66.7) |
| Mean *I*/σ(*I*) | 12.45 (2.00) | 9.8 (1.61) |
| Wilson B-factor | 25.4 | 60.31 |
| *R*<sub>merge</sub> | 0.119 (0.647) | 0.055 (1.13) |
| *R*<sub>meas</sub> | 0.129 (0.702) | 0.059 (1.23) |
| *R*<sub>pim</sub> | 0.049 (0.270) | 0.022 (0.487) |
| CC1/2 | n.a. (0.815) | 0.998 (0.549) |
| Resolution range for refinement (Å) | 49.55–2.08 (2.13–2.08) | 36.58–2.3 (2.38–2.3) |
| Reflections used in refinement | 43,887 (3,223) | 16,347 (665) |
| Completeness (%) | 99.4 (99.3) | 74.38 (30.42) |
| Reflections used for R-free | 2,297 (156) | 842 (45) |
| R-work | 0.178 (0.226) | 0.253 (0.371) |
| R-free | 0.233 (0.276) | 0.295 (0.425) |
| Number of non-hydrogen atoms | 5,868 | 2,878 |
| macromolecules | 5,747 | 2,867 |
| ligands | 8 | 1 |
| solvent | 113 | 10 |
| Protein residues | 705 | 360 |
| RMS(bonds) | 0.014 | 0.003 |
| RMS(angles) | 2.007 | 0.57 |
| Ramachandran favored (%) | 98.0 | 96.1 |
| Ramachandran allowed (%) | 2.0 | 3.9 |
| Ramachandran outliers (%) | 0.00 | 0.00 |
| Rotamer outliers (%) | 2.00 | 0.00 |
| Clash score | 5.0 | 5.40 |
| Average B-factor | 27.6 | 77.4 |
| macromolecules | 27 | 77.5 |
| ligands | 33.6 | 79.8 |
| solvent | 30.5 | 55.9 |

Statistics for the highest-resolution shell are shown in parentheses.

At the interface of the LysM and M23 domains, Domain 1 is predominantly negatively charged and Domain 3 is positively charged. At the center of the interface are van der Waals interactions between L109 and L344 in Domain 1 and 3, respectively. Additionally, the carboxyl group of D112 hydrogen-bonds with the hydroxyl group of Y330. Furthermore, Domain 1 residue E103 forms an electrostatic interaction with R371, present on the loop adjacent to the M23 catalytic groove. This combination of electrostatic and van der Waals interactions presumably serve to enhance interaction between the two domains and further appear to occlude the catalytic groove of the M23 domain. At the end of Domain 3, the ShyA polypeptide forms an amphipathic C-terminal α-helix where hydrophobic residues pack against the β-sheet of Domain 2 and hydrophilic residues are solvent exposed. A similar helix is also found in the Domain 3 of *Neisseria* structures for the EP NGO_1686 [PDB entries 6MUK and 3SLU; (47)], but not observed in the ShyB crystal structure because the protein construct did not include residues at the C-terminus. The presence of this helix in multiple structures may indicate a role in the orientation of Domain 2. The active site residues of ShyA’s M23 domain are similar to other M23 endopeptidases, in particular to ShyB and NGO_1686 (46, 47). The catalytic Zn^2+^ ion is ligated directly by H297, D301 and H378, and through water molecules to H376 and H345 (**Fig 2E**). Most of these residues are positioned on an antiparallel beta sheet in Domain 3. However, residue H297 is on an extended loop between two antiparallel beta strands.

### Structure of ShyA^OPEN^

Unexpectedly, the tertiary structure of ShyA obtained from the monoclinic crystals grown from the C-terminal His-tagged protein was drastically different from the ShyA^CLOSED^ structure (**Fig. 2B,C**). For reasons outlined below, we will refer to this structure as ShyA^OPEN^. Although each domain adopts a very similar conformation as the corresponding domain in ShyA^CLOSED^ (Cα r.m.s.d. range from 0.6 to 1.0 Å), the molecule is extended such that there are no interactions between individual domains within one molecule, except for the C-terminal helix, which still packs against Domain 2 as it does in ShyA^CLOSED^. Although diffraction data anisotropy resulted in an electron density map significantly less well-resolved than ShyA^CLOSED^, the electron density for the interdomain linkers in ShyA^OPEN^ was clearly resolved in the initial molecular replacement map and subsequent composite, simulated annealing omit maps of the entire structure (**Fig. S2B,C**). Because the interdomain linkages are likely flexible, an open form in solution could adopt many conformations and not only the overall shape observed in the crystal.

In order to transition between closed and open forms, the ShyA molecule must undergo a 138° rotation of Domain 2 around an axis passing through the Domain 2 – Domain 3 linker (near residue 265) (**Figure 2D**). Most of the changes occur in the interdomain linkages. Because of the rotation of Domain 2, the linker between Domain 3 and the C-terminal helix (∼396–400) also must change significantly (**Fig. S2C**). The major consequence of the observed conformational change is that Domain 1 has moved significantly away from Domain 3, making the presumed substrate binding site in Domain 3 completely solvent accessible.

We hypothesized how a minimal model of the D-Ala–DAP dipeptide crosslink (a part of the natural substrate for EPs) containing the scissile bond might be positioned in the putative substrate binding region of ShyA^OPEN^ (**Figure S2D).** Although not shown in the figure, the scissile amide bond carbonyl points towards the catalytic Zn [as proposed for the catalytic mechanism of the M23 peptidase LytM with transition state analog (50)]. In ShyA^CLOSED^, Domain 1 completely obscures the binding site region to the “right” of the Zn atom when viewed in the orientation of **Fig. S2E**. This modeling approach allowed us to test predictions about catalytic residues that are expected requirements for EP function. We created mutations in such residues and tested the resulting mutant protein’s activity by its ability to complement (or not) growth of a Δ6 endo strain. Specifically, we tested H345 (predicted to ligate the catalytic water), H297 (coordination of the catalytic zinc), R286 and D301 (predicted to be important based on its proximity to the active site zinc) (**Fig. 2E**). Consistent with important roles in zinc and water coordination, mutating R286, H297 or H345 to an alanine completely abrogated ShyA activity *in vivo* (**Fig. 2F**), while expression levels for these mutants were not affected (**Fig. S1C**). We had previously found that an H376A mutation rendered the protein non-functional as well (23), also consistent with its role in hydrogen-bonding to the Zn-bound catalytic water molecule. Thus, consistent with our structural prediction, residues D301, H345, H378, R286, H297 and H376 are crucial for ShyA’s EP function, and thus serve to functionally validate our structure’s *in vivo* significance.

In summary, our structural data suggest that ShyA likely assumes two conformations in equilibrium. The closed form is inactive and only the open form is predicted to be active due to PG substrate accessibility. We hypothesize that the *in vitro* activity we observe with purified ShyA is the result of conformational switching of a subfraction of ShyA molecules in solution. Indeed, we previously noticed that the C-terminally tagged construct that yielded the open conformation was highly active on its PG substrate (reaction times 30 min – 3 h) (23), while the N-terminally tagged construct (as well as untagged ShyA) required much longer reaction times of 12–16 hours (42) to complete PG digestion.

### Mutations predicted to favor the open conformation increase ShyA toxicity in vivo

Based on our structural predictions that allowed us to assign a putative inhibitory role for Domain 1, we created mutations that we expected to reduce interactions between Domain 1 and Domain 3 (**Fig. 3A**). We started by making targeted domain deletions and assessed their toxicity and function. We first created a truncated protein that lacks Domain 1 (Dom1) altogether, by fusing a standard periplasmic signal sequence (of DsbA (51)) with ShyA residues 161–430. To control for protein secretion effects, we also constructed a full-length ShyA version that had its native signal sequence (residues 1 - 35) replaced with that of DsbA (DsbA_ss_-ShyA). Overexpressing ShyA^ΔDom1^ construct in a wild-type background yielded no reduction in plating efficiency; however, colonies were generally smaller than the controls (**Fig. 3B**). Upon visualization of cells overexpressing ShyA^ΔDom1^, however, we noticed striking morphological defects. Most notably, we observed a high degree of sphere formation, indicative of increased PG cleavage activity (**Fig. 3C**). Overexpression of wild-type ShyA, an active site mutant (ShyA^H376A^), DsbA_ss_-ShyA or a Dom1+Dom2 truncation (residues 266–430) did not cause any growth or morphological defects (**Fig. 3B,C**). Thus, Domain 1 plays an important role in suppressing ShyA overexpression toxicity, consistent with a role in active site inhibition.

**Figure 3.**
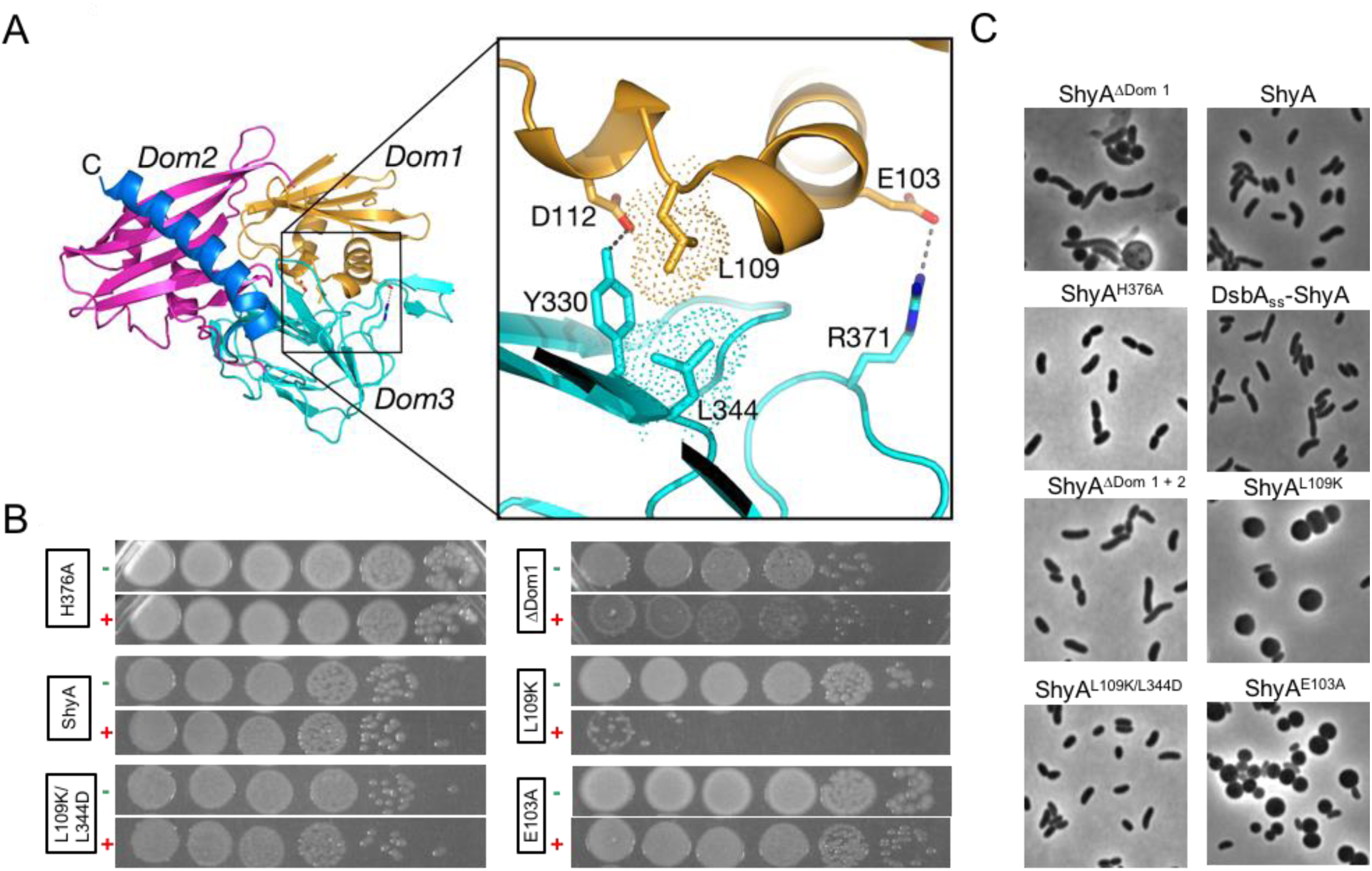
ShyA activity is enhanced by mutating Domain 1 – Domain 3 interface residues. **(A)** Overall domain structure of ShyA with zoomed-in view of residues predicted to mediate Domain 1 – Domain 3 interactions (insert). Domain colors are as in **Fig. 2B-D**. **(B)** Wild-type *V. cholerae* carrying the indicated mutant construct (arabinose-inducible) was diluted and spot-plated on plates containing either no inducer (–) or 0.2 % arabinose (+). **(C)** Wild-type *V. cholerae* carrying the indicated plasmid construct was grown to exponential phase (OD_600_∼0.4) in LB/chloramphenicol (20 µg/mL) and induced for 3 h with 0.2 % arabinose, followed by imaging on an agarose pad.

We next focused our attention on the hydrophobic interactions between L109 and L344, the salt bridges between D112/Y330 and E103/R371, and the general acidic-basic interaction of the entire domain interface (**Fig. 3A**) to create additional mutants that might favor an open conformation. We predicted that mutating L109 to a basic residue would strongly repel the basic patch in Domain 3 and also sterically clash with L344 in Domain 3, opening the structure. Remarkably, and consistent with this hypothesis, overexpression of the ShyA^L109K^ mutant caused a near 100,000-fold plating defect and resulted in population-wide sphere formation when expressed in liquid medium (**Fig. 3B,C**). To further probe the L109-L344 interaction’s involvement in Domain 1 – 3 interactions, we reasoned that a negative charge addition in the opposing L344 residue would electrostatically interact with the K109 residue to promote the closed conformation and reduce ShyA^L109K^ toxicity. Consistent with a functional interaction between L109 and L344, mutating L344 to an aspartic acid (L344D) in the L109K background restored wild-type growth on a plate and normal morphology (**Fig. 3B,C**).

We next turned to the predicted salt bridge between E103 and R371. To potentially interrupt this interaction and consequently destabilize Domain 1 – Domain 3 interactions, we introduced an alanine substitution in E103. The resulting ShyA^E103A^ mutant caused loss of rod shape and sphere formation, yet no plating defect, similar to the ΔDom1 mutant (**Fig. 3C**). Western Blot analysis revealed that L109K and E103A were expressed at levels similar to wild-type ShyA (**Fig. S1**), suggesting that mutant toxicity was not simply the result of enhanced protein accumulation. Lastly, we also tested mutations in D112 (creating ShyA^D112E^ and ShyA^D112A^). D112 is predicted to hydrogen bond with Y330 in Domain 3. Again, consistent with disruption of this stabilizing interaction, both ShyA^D112E^ and ShyA^D112A^ caused morphological defects upon overexpression as well as a mild plating defect (small colonies) (**Fig. S3**). Taking all *in vivo* results together, we conclude that mutations that are expected to disrupt interactions between Domains 1 and 3 activate the protein to varying degrees.

### Mutations predicted to favor the open conformation increase ShyA activity in vitro

While predicted open conformation mutants exhibited higher toxicity *in vivo,* we sought to confirm that this was indeed due to higher PG cleavage activity. We purified N-terminally tagged ShyA^67-426^ as well as its derivatives ShyA^L109K^ and ShyA^L109K/L344D^ and measured their activity by assaying digestions of purified whole PG sacculi. With ShyA^67-426^, we observed slow digestions proceeding to completion after 20 hours, but comparatively little activity after short (10 min – 3hours) incubation periods (**Fig. 4A, Fig. S4**). Thus, this version of ShyA (retaining domains in the ShyA^CLOSED^ structure) is active at a low level. In contrast, the ShyA^L109K^ derivative of this construct almost completely digested purified sacculi within 10 minutes with no further activity after 20 hours of incubation (**Fig. 4B**). Thus, the L109K mutation enhances PG cleavage activity *in vitro*. ShyA^L109K/L344D^ exhibited significantly reduced activity compared to ShyA^L109K^ at the earliest time point (**Fig. 4B**), though activity was still considerably higher than that of wild-type ShyA and remained at ShyA^L109K^ levels for the remainder of the time course. Thus, establishing an ionic bridge between Domains1 and 3, while drastically reducing toxicity *in vivo*, still results in a considerably faster reaction rate than the wild-type protein. We speculate that the difference between *in vivo* and *in vitro* results reflects that the geometry of the engineered ionic bridge is not ideal and the two residues may still sterically clash *in vitro*. The specific periplasmic environment might reduce the conformational consequences of this steric clash, for example through molecular crowding, a condition not readily reproducible *in vitro*. Another observation lent support to the idea that the ShyA^L109K/L344D^ has reduced activity *in vivo*; we found that this variant only partially complemented a Δ6 endo strain (**Fig. S5**). However, both *in vitro* and *in vivo*, the L344D mutation reduced activity of ShyA^L109K^, providing additional evidence that enhancing Domain1 – Domain3 interactions renders ShyA less active than reducing such interactions.

**Figure 4.**
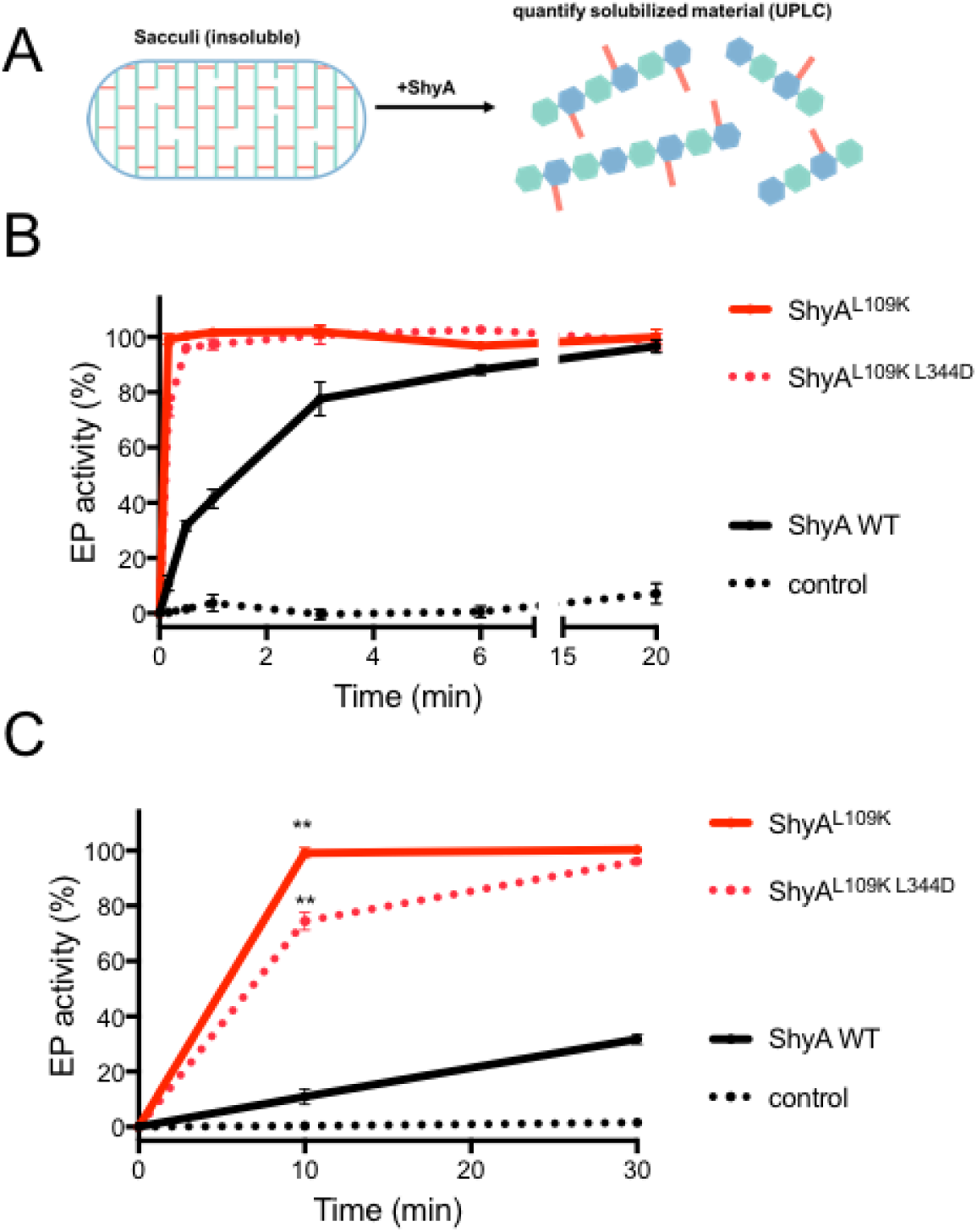
*In vitro* toxicity of hyperactive ShyA variants. **(A)** Overview of reaction. Purified (insoluble) PG sacculi are digested with ShyA and the solubilized material is quantified via UPLC. **(B)** Purified (N-terminally His-tagged) ShyA and its derivatives were mixed with purified PG sacculi and incubated at 37°C. At the indicated time points, samples were withdrawn, reactions stopped by boiling, and digested with muramidase, followed by analysis on UPLC. Activity was calculated as area under the curve of both solubilized PG material and remaining pellet (see Methods for details). Data are averages of three independent reactions; error bars represent standard deviation. **(C)** Enhanced (first 30 min reaction time) view of the graph in **(B)** to illustrate significant difference between ShyA^L109K^ and ShyA^L109K/L344D^. ** indicates statistical significance (t-test, p<0.001). The control is purified PG without addition of EPs.

### Conformational regulation may be widespread among divergent M23 endopeptidases

We asked whether M23 EPs from other bacteria might exhibit a mode of regulation similar to the one we are proposing for ShyA. Indeed, published structures of EPs from divergent pathogens, namely Csd3 (*Helicobacter pylori*) and NGO_1686 (*Neisseria gonorrheae*) appeared to assume a domain organization remarkably similar to that of ShyA (**Fig. 5A**, (47, 48)). In addition, a homology model was created for MepM (*E. coli*) using the ShyA^CLOSED^ structure. We created mutations in NGO_1686 of *Neisseria gonorrheae* and MepM of *E. coli* to test predictions about enzyme activation. For NGO_1686, we identified a residue (E132) that we implicated in inter-domain connections (via a salt-bridge between E132 and R293) (**Fig. S6**) and tested a mutation (E132R) that we predicted to destabilize this interaction, analogous to E103A in ShyA. The MepM model does not have a clearly identifiable acidic-basic Domain 1 – Domain 3 interface, and we thus simply created a ΔDomain1 variant that should relieve catalytic inhibition by Dom1.

**Figure 5.**
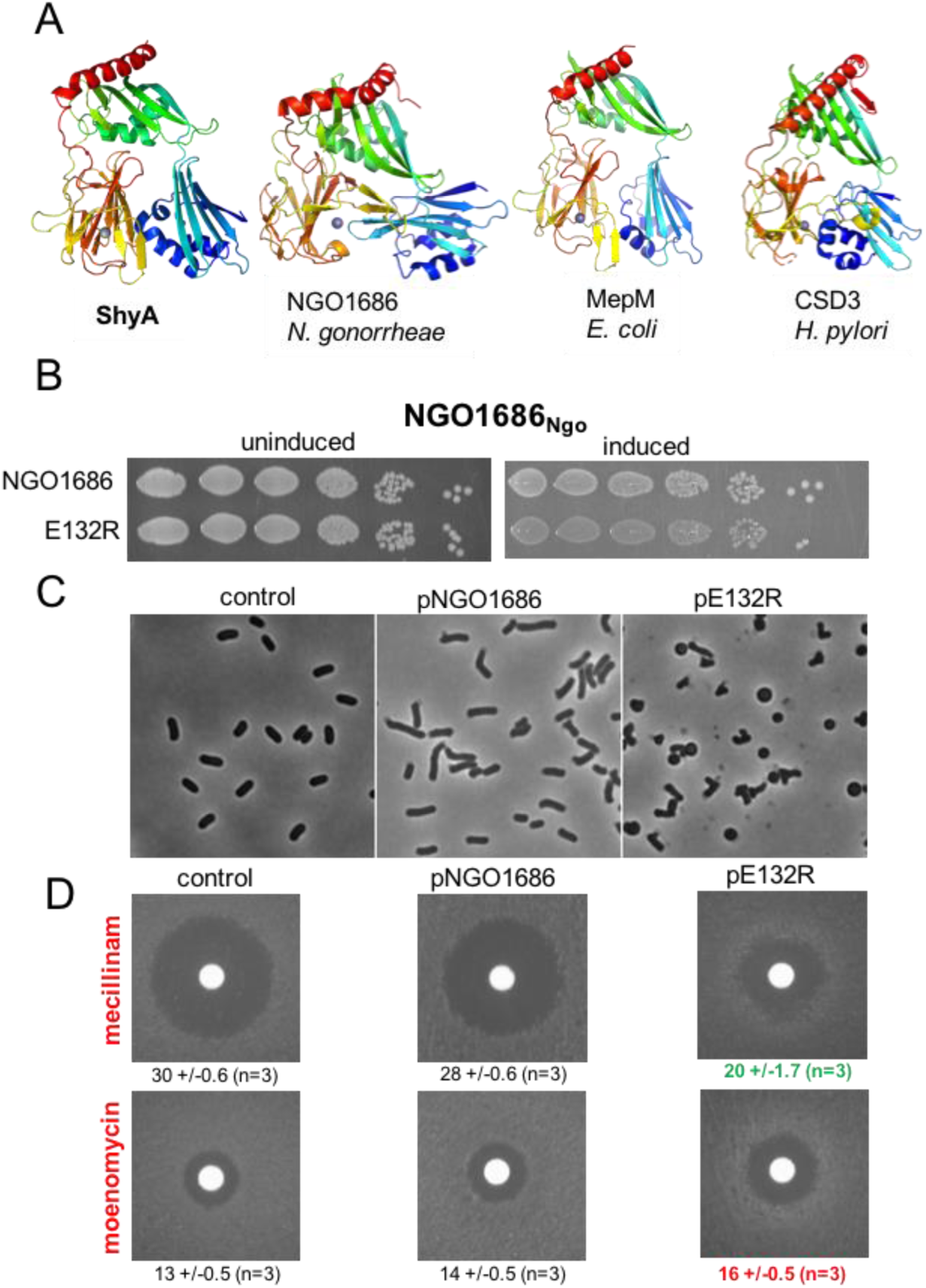
ShyA functional domain organization is conserved in divergent Gram-negative pathogens. **(A)** Domain organization of LysM/M23 EPs. Structures were obtained either from published sources (CSD3 and NGO1686, PDB entries 4RNY and 6MUK, respectively) or modeled onto our ShyA structure using the Swiss Model server (71) (MepM). Polypeptides are rainbow colored from blue at N-terminus to red at C-terminus **(B)** Overexpression toxicity of NGO1686^E132R^. Strains carrying arabinose-inducible EPs were plated in the absence (uninduced) or presence (induced) of 0.2 % arabinose. **(C)** MG1655 carrying pBAD33 or its derivatives containing wild-type NGO1686 or NGO1686^E132R^ were grown to mid-exponential phase (OD_600_ ∼ 0.4) and induced with 0.2 % arabinose, followed by imaging on agarose pads. **(D)** MG1655 carrying the indicated plasmids was plated evenly (∼10^8^ cfu) on LB agar containing 0.2 % arabinose and chloramphenicol (20 µg/mL). A filter disk containing mecillinam (10 µL of 20 mg/ml stock) or moenomycin (10 µL of 5 mg/mL stock) was then placed in the center of the plate, followed by incubation for 24 hours at 37°C. Numbers below the images indicate zone of inhibition in mm +/– standard deviation. Control = ShyA^H376A^, Eco, *E. coli,* Ngo, *Neisseria gonorrheae*.

We expressed NGO_1686 and its E132R variant heterologously in *E. coli.* While neither the mutant variant nor the wild-type protein caused a strong plating defect upon induction (although colonies were smaller for the mutant form, **Fig. 5B**), expression of NGO1686^E132R^ caused a dramatic morphological defect, including sphere formation (**Fig. 5C**). A slight defect was visible even in the wild-type protein, but this was exacerbated in the mutant. To assess enhanced NGO_1686 toxicity more quantitatively, we also performed a mecillinam/moenomycin sensitivity assay. In *E. coli*, increased EP activity imparts mecillinam resistance, likely due to overactivation of aPBPs (52). We predicted that such aPBP overactivation should also render these cells more susceptible to moenomycin, an aPBP inhibitor (53). We thus performed a Kirby-Bauer zone of inhibition assay (54) on *E. coli* overexpressing an inactive version of ShyA (ShyA^H376A^, negative control), NGO_1686 or its E132R derivative. Overexpression of the NGO1686^E132R^, but not the wild-type protein or the inactive ShyA^H376A^, drastically increased resistance against mecillinam and sensitivity to moenomycin (**Fig. 5D**), again suggesting that indeed the E132R mutation enhances NGO_1686 activity.

Overexpressing DsbAss-MepM^ΔDom1^ in its native host *E. coli* dramatically reduced plating efficiency compared to the wild-type version of the protein (**Fig. S7**). Interestingly, DsbAss-MepM (the true parental to MepM^ΔDom1^) exhibited slightly enhanced toxicity (manifesting as small colonies) compared to the wild-type protein, suggesting that soluble MepM by itself is more active than its predicted membrane-bound form. In cumulation, we demonstrate that ShyA’s domain organization is conserved across divergent Gram-negative pathogens and suggest that the inhibitory role of Domain 1 constitutes a widespread mechanism of regulation.

## Discussion

The mechanism of regulation of endopeptidases during cell elongation is poorly understood for Gram-negative bacteria. Data from *E. coli* and *P. aeruginosa* point towards proteolytic turnover (44, 45); however, in *E. coli* this mechanism of regulation appears to be more dominant during stationary phase remodeling and its role in *P. aeruginosa* is unclear. Notably, deleting EP-specific proteases only causes minor phenotypes under laboratory growth conditions and does not seem to affect cell elongation in exponential phase unless severe osmotic conditions are applied (44, 45). Here, we present data suggesting a conformational switch mechanism of activation for LysM/M23 EPs to effectively regulate cell wall remodeling during cellular expansion. Interestingly, ShyA mutants that we predicted to favor active conformations differed in the severity of their toxicity. For example, only the L109K mutation caused a strong plating defect upon induction. The ΔDom1, E103A and D112E mutations caused sphere formation similar to the L109K mutation; however, these former mutants were able to ultimately form colonies on a plate. The open conformation observed in the crystal structure likely represents an extreme (possibly induced by high protein concentrations that promoted intermolecular Dom1-Dom3 interactions, stabilizing the wide-open conformation), while the situation *in vivo* is likely more dynamic, with various degrees of opening up of the ShyA^CLOSED^ structure. It is possible that the L109K change simply results in the strongest repellent effect between Dom1 and Dom3, while E103A/D112E present as a more dynamic form with an open-closed equilibrium that is just pushed slightly towards the open state.

Our data raise the possibility that bacteria maintain a large pool of inactive EP precursors that are actuated only when or where needed. This strategy, as opposed to transcriptional regulation, would allow for quick responses to environmental conditions that may require enhanced cleavage activity. A large, inactive pool of EPs would also explain an apparent oddity. A modeling approach has suggested that less than 100 “dislocation events” (*i.e.*, PG cleavage events) per cell are sufficient for growth (55), and previous biochemical work demonstrated that a similarly low number of cell wall synthesis complexes is active per generation (56). Compared to these estimates, EP numbers have been determined to be much higher using ribosome profiling in the model organism *E. coli* (between ∼300 [MepM] and ∼4000 [spr] (57)). Our estimate for the number of ShyA molecules/cell (∼1500) is also within this range. Thus, the number of EP molecules in the cell vastly exceeds the number of active molecules theoretically required to sustain growth. It is tempting to speculate that these surplus molecules indeed provide a reservoir for changing growth conditions.

While we have not identified the signal promoting this conformational switch, we hypothesize that this might be an interaction with cell wall synthases or the protein complexes they are embedded in, to enable tight coordination between synthesis and degradation. Importantly, however, the aPBPs themselves are not likely activators of conformational switching – data in *E. coli* suggests that EPs are limiting for aPBP activity (52); if aPBPs were necessary to activate EPs, this relationship would be reversed. In *E. coli*, the lipoprotein NlpI was recently identified as a multifunctional regulator of EP activity and coordinator of hydrolysis with PG synthesis functions (40). NlpI was shown to inhibit the activity of the ShyA homolog MepM, and its *V. cholerae* homolog is thus likely not the ShyA activator. However, it is possible that similar protein-protein interactions may induce the observed conformational switch in EPs, analogous to activation of amidases by their EP-like regulators (14, 29, 58). Alternatively, EPs might recognize intrinsic substrate cues in PG, *e.g*., tension state, in line with Arthur Koch’s “smart autolysin” hypothesis (59). ShyA possesses a LysM carbohydrate binding domain (49) in Domain 1, and in the smart autolysin scenario, an interaction of this domain with PG might enhance the switch to an active form under specific (e.g., osmotic challenge) conditions. From a broader perspective, our data add to the emerging theme of a multifaceted regulatory system for EPs (transcriptional regulation, post-translational degradation, conformational change). This complex system likely reflects the necessity of tightly controlling cell wall degradation during cell elongation in rapidly changing environments.

## Materials and Methods

### Media, Chemicals and Growth conditions

Strains were grown in LB medium (Fisher Scientific, cat # BP1426-2) at 37°C, shaking at 200 rpm, unless otherwise indicated. Antibiotics were used at the following concentrations: streptomycin (200 µg/mL), chloramphenicol (20 µg/mL), kanamycin (50 µg/mL). Inducers were used at 0.2 % (arabinose) or 300 µM (IPTG), respectively.

### Bacterial Strains and Plasmids

Bacterial strains and oligos are summarized in **Tables S1 and S2**. All *V. cholerae* strains are derivatives of N16961 (60). Subcloning was conducted in *E. coli* SM10 lambda pir. Isothermal assembly (ITA, Gibson Assembly (61)), was used for all cloning procedures. ShyA and ShyC were cloned into pBADmob (a pBAD33 derivative containing the mobility region from pCVD442 (62)) using primer pairs TDP484/1302 (ShyA) or TDP486/779 (ShyC).

Site-directed mutagenesis was performed using the NEB Q5 site-directed mutagenesis kit (NEB, Ipswitch, MA, cat #E0554S) using mutagenesis primers summarized in **Table S2** with pBADmob*shyA* as a template.

pBADmob*NGO_1686* was constructed by amplifying the *ngo1686* orf from heat-lysed *Neisseria gonorrhea* ATCC 49226 with primers TDP1365/1367, followed by cloning into Sma1-digested pBADmob via ITA. This plasmid was then used as a template for construction of E132R using the NEB Q5 site-directed mutagenesis kit.

### Expression and Purification of Recombinant ShyA with N-terminal His-tag

After predicting the periplasmic signal sequence of ShyA with SignalP 4.1, the coding region (residues 36–430 for t-ShyA or residues 67–426 for a shorter truncated s-ShyA) of the ShyA gene was amplified from *V. cholerae* N16961 genomic DNA using mutagenic primers TD-JHS062 and TD-JHS063 (for t-ShyA) and TD-JHS323 and TD-JHS324 (for s-ShyA). The PCR products were digested with NdeI and BamHI and inserted into the pET-15b plasmid (Novagen) digested with the same enzymes. To create ShyA mutant proteins, site-specific mutagenesis primers (L109K and L109K/L344D, designed with the NEBaseChanger mutagenesis tool) were used with the NEB Q5 site-directed mutagenesis kit (New England Biolabs).

For the purification of 6xHis+ShyA proteins, *E. coli* BL21(DE3) (Novagen) was transformed with the resulting recombinant plasmids [pET15bt-ShyA, pET15bs-ShyA, pET15bs-ShyA(L109K), pET15bs-ShyA(L109KL344D)]. An overnight culture from a single colony was used to inoculate 1 liter of Luria-Bertani medium. Cells were grown with vigorous shaking (220 rpm) at 37°C to an optical density at 600 nm (OD_600_) of 0.5 and were induced with 1 mM (final concentration) isopropyl-β-D-thiogalactopyranoside (IPTG, Gold Bio, cat# I2481C25) for 12 h at 30°C. Harvested cells were re-suspended in binding buffer JHS1 [20 mM Tris-HCl (pH 7.9), 0.5 M NaCl, and 25 mM imidazole] and lysed by sonication. N-terminally tagged ShyA and its variants proved insoluble, so inclusion bodies were isolated after centrifugation at 21,002 rcf for 45 min. For resolubilization of insoluble ShyA from inclusion bodies, pellets were re-suspended in solubilization buffer JHS4 [20 mM Tris-HCl (pH 7.9), 0.5 M NaCl, 25 mM imidazole, and 3 M urea] and incubated overnight on a rotator at 4°C. Supernatant containing resolubilized ShyA was centrifuged at 21,002 rcf for 45 min and loaded onto HisPur^TM^ Cobalt column(Thermo Scientific, cat# 89964), which was then washed with 6 volumes of solubilization buffer JHS4 followed by 6 volumes of washing buffer JHS5 [20 mM Tris-HCl (pH 7.9), 0.5 M NaCl, 50 mM imidazole, and 3 M urea]. ShyA protein was eluted with 10 volumes of elution buffer JHS6 [20 mM Tris-HCl (pH 7.9), 0.5 M NaCl, and 3 M urea] containing a linear imidazole gradient from 100 to 500 mM. Fractions containing ShyA protein were pooled and dialyzed against JHS-D1 buffer [20 mM Tris-HCl (pH 7.9), 250 mM NaCl, 10 % (v/v) glycerol, 2 mM EDTA, and 0.5 M urea] to remove imidazole and cobalt ions, then dialyzed against buffer JHS-D2 [20 mM Tris-HCl (pH 7.9), 150 mM NaCl, 20 % glycerol, and 0.5 mM ZnSO_4_] to remove urea and add back zinc, and finally buffer JHS-D3 [20 mM Tris-HCl (pH 7.9), 150 mM NaCl, and 30 % glycerol].

### Crystallization and structure solution of ShyA with N-terminal His6 tag

6xHis+t-ShyA was purified as above and dialyzed in JHS-FPLC buffer [20 mM Tris-HCl (pH 7.9) and 150 mM NaCl], concentrated by centrifugal filter devices (Millipore, 10,000 MW CO), and then further purified by FPLC size exclusion chromatography using a HiLoad 16/60 Superdex 200 prep grade column (GE Healthcare Life Sciences). Pure 6xHis+t-ShyA fractions were collected and concentrated up to 11.4 mg/ml. Protein concentration was measured via UV absorbance at 280 nm, and calculated using ShyA’s molar extinction coefficient (ε_280_ = 37550 M^−1^ cm^−1^, http://expasy.org/cgi-bin/protparam). Protein purity was confirmed by SDS-PAGE with Coomassie blue staining.

For crystallization of ShyA, the concentration of FPLC-purified ShyA was adjusted to 2.7 mg/mL. Concentrated protein was combined with purified peptidoglycan sacculi (prepared as described previously (23)) and crystallization conditions were screened at room temperature using a Crystal Phoenix liquid handling robot (Art Robbins Instruments). Crystals optimally grew by mixing protein and precipitant (0.2 M malonic acid pH 6.0 and 16% PEG 3350) and incubating by hanging drop vapor diffusion. Rectangular cuboid crystals formed from these conditions after 3 days at room temperature. Crystals were then transferred to cryoprotectant consisting of precipitant containing 25% (v/v) glycerol. Crystals were frozen with a liquid nitrogen stream at MacCHESS beamline F1 at the Cornell High Energy Synchrotron Source. The diffraction data were indexed and scaled using the HKL2000 software (63). The crystal structure was solved by molecular replacement using the AMoRe Suite (64) package in CCP4 (Collaborative Computational Project, 1994) with ShyB (PDB entry 2GU1) as a search model. The ShyA^CLOSED^ model was built using iterative cycles of model building with COOT software (65) and Refmac5 (66) for refinement. Data and refinement statistics are shown in **Table 1**. ShyA^CLOSED^ was deposited in the PDB under entry 6UE4.

### Purification, crystallization, and structure solution of ShyA with C-terminal His-tag

Competent *E. coli* Rosetta-gami 2 (DE3) (Novagen) cells were transformed with pET28-VcShyA(36–430)-His6 (23) in the presence of chloramphenicol (Cm, 100 μg/ml) and kanamycin (Kan, 40 μg/ml). Two 2.8-L Fernbach flasks each containing 450-ml Terrific Broth (Research Products International) were inoculated 1:100 with an overnight culture and grown in the presence of Cm and Kan at 37° with shaking. When the cultures reached an optical density (at 600 nm) of 1.0 (∼7 hours), flasks were transferred to a precooled 22°C incubator for one hour before inducing each with 0.4 mM IPTG and shaking for another 17 hr at 22°C. After centrifugation, the cell pellet from each flask was suspended in 30 ml lysis buffer (300 mM NaCl, 20 mM Na phosphate pH 7.5) containing one-half of a Roche cOmplete™ Protease Inhibitor Cocktail (EDTA-free) tablet and frozen at –80°C.

After thawing the cell suspension from one flask, the other half of the protease inhibitor tablet was added to the suspension along with 5 μg/ml DNase I. The suspension was passed three times through an Emulsiflex-C3 homogenizer at 4°C (air pressure ∼20,000 PSI), and then centrifuged in a Sorvall SS-34 rotor at 20,000 rpm (48,000*g*) for 30 minutes. Finally, the lysate was filtered through a 0.45 μ filter.

A 6.5 ml Tricorn column (GE Healthcare) was packed with HisPur™ Cobalt Resin (Thermo Fisher) and preequilibrated with buffer A (300 mM NaCl, 50 mM Tris-HCl pH 7.5). The lysate was loaded onto the column at 1 ml/min, washed at 2 ml/min with buffer A, and then washed with 2% and 4% buffer B (300 mM NaCl, 300 mM imidazole, 50 mM Tris-HCl pH 7.5) until absorbance at 280 nm reached base line. The column was eluted with 60% buffer B until *A*_280_ reached base line. Protein was concentrated to 20 mg/ml, and 10% (v/v) glycerol was added. Aliquots of 0.5 ml were frozen in liquid nitrogen and stored at –80°C.

Before setup of crystallization experiments, a frozen protein aliquot was thawed and loaded onto a Superdex HR 75 10/300 (GE Healthcare) size exclusion column and eluted with 150 mM NaCl, 20 mM Tris-HCl, pH 7.6 at 0.5 ml/min. The protein peak was analyzed with SDS-PAGE and the purest fractions were pooled and concentrated to 17.5 mg/ml with an Amicon Ultra-4 30K MWCO concentrator (EMD-Millipore). Protein was screened for crystallization with the Qiagen commercial kits PEG Suite and Classics in 96-well Intelli-Plates (Art Robbins). Sitting drops containing 0.5 μl protein and 0.5 μl precipitant, were equilibrated against 100 μl precipitant in the reservoir. The crystal chosen for data collection grew in precipitant containing 0.25 M sodium citrate, 10% polyethylene glycol 3350, 0.1 M Tris-HCl pH 6.0. Crystals were transferred to a cryosolvent consisting of 10% (w/v) ethylene glycol in the precipitant, and frozen in liquid nitrogen.

Diffraction data from monoclinic crystals obtained with protein containing the C-terminal His6 tag were measured at the LS-CAT beamline 21-ID-D, Advanced Photon Source at Argonne National Laboratory, equipped with a Dectris Eiger 9M detector running in continuous mode. Eighteen hundred images, 0.2° oscillation angle each, were processed with autoPROC (Global Phasing Limited) (67). Data collection statistics are summarized in **Table 1**. Due to significant anisotropy in diffraction limits, an ellipsoidal region of reciprocal surface was calculated by STARANISO in autoPROC and used as a cutoff. Only reflections within this ellipsoid were used for the structure refinement. Anisotropic Debye-Waller corrections were also applied.

Molecular replacement calculations with Phaser (68) failed to produce a solution when either ShyB [PDB id 2GU1 (46); 50% sequence identity to ShyA] or the ShyA^CLOSED^ structure was used as a search model. Only after dividing the latter model into individual domains as search models was a significant solution obtained with a log likelihood gain of 1,840 and translation function Z-score of 41.5. The electron density map was interpretable with the aid of the individual domain structures from the ShyA^CLOSED^ model. Interdomain linkers were clearly resolved confirming the unique structure observed (**Figure S2**). Electron density for the C-terminal α-helix could be interpreted with the help of the ShyA^CLOSED^ structure. The model was built with Coot (65) and refined with Phenix (69) with data to 2.3-Å resolution to a *R*work/*R*free of 0.25/0.29 with excellent geometry. The structure, referred to here as ShyA^OPEN^, was deposited in the PDB as entry 6U2A.

### Western Blotting

Whole cell extracts (30 µg) were resolved by 10% SDS-PAGE and the proteins were transferred to a membrane by using a semi-drying transfer system (iBlot 2, Invitrogen). The membrane was then blocked with blocking solution containing 4% milk (dry skim milk dissolved in 20 mM Tris-HCl (pH 7.8), 150 mM NaCl, 0.1% Triton X-100) overnight. The membrane was incubated with anti-ShyA polyclonal antibody (1:5,000, produced by Pocono Rabbit Farm & Laboratory, PA) or monoclonal anti-αRpol antibody (1:15,000, BioLegend cat# 663104) for two hours and then washed with TBST (20 mM Tris-HCl (pH7.8), 150 mM NaCl, 0.1% Triton X-100). The washed membranes were further incubated with anti-rabbit (1:15,000, Li-Cor cat# 926-32211) or anti-mouse secondary antibodies (1:15,000, Li-Cor cat# 926-32210) for 1 hour. All blots were washed three times with TBST, scanned on an Odyssey CLx imaging device (LI-COR Biosciences), and visualized using Image Studio™ Lite (Li-Cor) software.

### Endopeptidase activity assays

*In vitro* assays to test the degrading activity of the different ShyA variants were performed on stationary phase *Vibrio cholerae* sacculi (70). Ten micrograms of substrate were incubated with 10 µg of every enzyme (or H_2_O in the negative control) in 50 mM TrisHCl pH 7.5, 100 mM NaCl buffer (total reaction volume 50 μl) during a time course experiment (10 min, 30 min, 1 h, 3 h, 6 h and 20 h). Prior to the muramidase digestion, the individual enzymatic reactions were heat-inactivated for 10 min at 100 °C. Soluble (released muropeptides and fragments) and pellet (intact sacculi) fractions were separated by centrifugation at 22,000 x g for 15 min. Pellet fractions were resuspended in 40 µl of water. Two microliters of muramidase (1 mg/ml) were added to both fractions, and samples were further incubated at 37 °C for 16 h. Reactions were heat inactivated for 10 min at 100 °C, soluble products reduced with sodium borohydride, and their pH adjusted.

The samples were injected into a Waters UPLC system (Waters, Massachusetts, USA) equipped with an ACQUITY UPLC BEH C18 Column, 130 Å, 1.7 μm, 2.1 mm × 150 mm (Waters) and a dual wavelength absorbance detector. Eluted fragments were separated at 45°C using a linear gradient from buffer A [formic acid 0.1% (v/v)] to buffer B [formic acid 0.1% (v/v), acetonitrile 40% (v/v)] in an 18 min run with a 0.250 ml/min flow, and detected at 204 nm. Identification of muropeptides was performed by mass spectrometry following the same separation methods described above on a UPLC system coupled to a Xevo G2-XS QTof quadrupole time-of-flight mass spectrometer (Waters MS Technologies). Relative activity was calculated as the percent of muropeptides released compared to the ShyA WT activity at 20 h and the decrease in substrate amounts, over a time course experiment performed in triplicates. Statistical analysis was performed using multi-t-test in Prism Software (Graphpad Software, San Diego, CA).

## Acknowledgements

We thank David Erickson (Cornell University) for providing us with *Neisseria gonorrhea*. Research in the Dörr laboratory is supported by National Institutes of Health (NIH) grants R01AI143704 and R01GM130971. Research in the Saper laboratory was funded by the NIH (R01GM130971) and the Department of Biological Chemistry, University of Michigan. Research in the Cava lab is supported by MIMS, the Knut and Alice Wallenberg Foundation (KAW), the Swedish Research Council and the Kempe Foundation. Work in the Mao laboratory was supported by NIH grant 5R01GM116964. This work is partially based upon research conducted at the Cornell High Energy Synchrotron Source (CHESS), which is supported by the National Science Foundation and the National Institutes of Health/National Institute of General Medical Sciences under NSF award DMR-1829070, using the Macromolecular Diffraction at CHESS (MacCHESS) facility, which is supported by award GM-124166 from the National Institutes of Health, through its National Institute of General Medical Sciences. This research used resources of the Advanced Photon Source, a U.S. Department of Energy (DOE) Office of Science User Facility operated for the DOE Office of Science by Argonne National Laboratory under Contract No. DE-AC02-06CH11357. Use of the LS-CAT Sector 21 was supported by the Michigan Economic Development Corporation and the Michigan Technology Tri-Corridor (Grant 085P1000817).

---

**Supplementary Table 1.** Strains and plasmids used in this study.

| Strain /plasmid |  | Relevant description | Reference /source |
| --- | --- | --- | --- |
| ***E. coli* strains** |  |  |  |
| DH5α λpir | DJ-ED1 | F- endA1 glnV44 thi-1 recA1 relA1 gyrA96 deoR nupG Φ80d/lacZΔM15 Δ(lacZYA-argF)U169, hsdR17(rK- mK+), λpir | Lab stock |
|  | DJ-ED2 | pET15b | This study |
|  | DJ-ED100 | pET15bt-ShyA | This study |
|  | DJ-ED101 | pET15bt-ShyA(H375A) | This study |
|  | DJ-ED102 | pET15bs-ShyA | This study |
|  | DJ-ED103 | pET15bs-ShyA(H375A) | This study |
|  | DJ-ED104 | pET15bs-ShyA(L109K) | This study |
|  | DJ-ED105 | pET15bs-ShyA(L109KL344D) | This study |
|  | DJ-ED106 | pET15bs-ShyA(Q193CN337C) | This study |
|  | DJ-ED107 | pET15bs-ShyA(S262CA398C) | This study |
|  | DJ-ED108 | pET28a6xHisSumo+s-ShyA | This study |
|  | DJ-ED109 | pET28a6xHisSumo+s-ShyA(H375A) | This study |
|  | DJ-ED110 | pET28a6xHisSumo+s-ShyA(L109K) | This study |
|  | DJ-ED111 | pET28a6xHisSumo+s-ShyA(L109KL344D) | This study |
|  | DJ-ED112 | pET28a6xHisSumo+s-ShyA(Q193CN337C) | This study |
|  | DJ-ED113 | pET28a6xHisSumo+s-ShyA(S262CA398C) | This study |
| BL21(DE3/pLysS) | DJ-EB1 | *B F- ompT gal dcm lon hsdSB(rB-mB-) λ(DE3 [lacI lacUV5-T7p07 ind1 sam7 nin5]) [malB+]K-12(λS) pLysS[T7p20 orip15A](CmR)* | Lab stock |
|  | DJ-EB2 | pET15b | This study |
|  | DJ-EB100 | pET15bt-ShyA | This study |
|  | DJ-EB101 | pET15bt-ShyA(H375A) | This study |
|  | DJ-EB102 | pET15bs-ShyA | This study |
|  | DJ-EB103 | pET15bs-ShyA(H375A) | This study |
|  | DJ-EB104 | pET15bs-ShyA(L109K) | This study |
|  | DJ-EB105 | pET15bs-ShyA(L109KL344D) | This study |
|  | DJ-EB106 | pET15bs-ShyA(Q193CN337C) | This study |
|  | DJ-EB107 | pET15bs-ShyA(S262CA398C) | This study |
| BL21(DE3)<br>RosettaGami 2 | DJ-ER200 | $\Delta(ara-leu)7697 \Delta lacX74 \Delta phoA$<br>*PvuII phoR araD139 ahpC galE galK*<br>*rpsL*(DE3) F'[*lac<sup>+</sup> lacI<sup>q</sup> pro*] *gor522::Tn10*<br>*trxB* pRARE2 (Cam<sup>R</sup>, Str<sup>R</sup>, Tet<sup>R</sup>) | Lab stock |
|  | DJ-ER201 | pET28a6xHisSumo | This study |
|  | DJ-ER202 | pET28a6xHisSumo+s-ShyA | This study |
|  | DJ-ER203 | pET28a6xHisSumo+s-ShyA(H375A) | This study |
|  | DJ-ER204 | pET28a6xHisSumo+s-ShyA(L109K) | This study |
|  | DJ-ER205 | pET28a6xHisSumo+s-ShyA(L109KL344D) | This study |
|  | DJ-ER206 | pET28a6xHisSumo+s-ShyA(Q193CN337C) | This study |
|  | DJ-ER207 | pET28a6xHisSumo+s-ShyA(S262CA398C) | This study |
| MG1655 | EP17 | MG1655 pBADmobngo_1686 | This study |
|  | EP18 | MG1655 pBADmobngo_1686 *E131R* | This study |
|  | EP19 | MG1655 pBADmobmepM | This study |
|  | EP20 | MG1655 pBADmobdsbAss-mepM | This study |
| | EP21 | MG1655 pBADmobdsbAss-mepM $\Delta Dom1$ | This study |
| **Plasmids** |  |  |  |
| pET15b | pDJ1 | Over expression vector in *E.coli* | Novagen |
| pET28a6xHisSumo | pDJ2 | Over expression vector in *E.coli* | Novagen |
| ***V. cholerae* strains** |  |  |  |
| N16961 | wild type | smR | Lab stock |
|  | EP1 | N16961 pBADmobshyA | This study |
|  | EP2 | N16961 pBADmobshyC | This study |
| | EP3 | $\Delta 6$ endo (N16961 $\Delta shyABC$ $\Delta vc1537$ $\Delta vca1043$ $\Delta vc0843$ $lacZ::P_{iptg:shyA}$) | This study |
| | EP4 | $\Delta 6$ endo $\Delta ldtA$ $\Delta ldtB$ | This study |
| | EP5 | N16961 pBADmobshyA $\Delta dom$ 1 | This study |
|  | EP6 | N16961 pBADmobshyA L109K | This study |
|  | EP7 | N16961 pBADmobshyA E103A | This study |
|  | EP8 | N16961 pBADmobshyA D112A | This study |
|  | EP9 | N16961 pBADmobshyA L109K L344D | This study |
|  | EP10 | EP3 pBADmobshyA | This study |
|  | EP11 | EP3 pBADmobshyA L109K L344D | This study |
|  | EP12 | EP3 pBADmobshyA R286A | This study |
|  | EP13 | EP3 pBADmobshyA H297A | This study |
|  | EP14 | EP3 pBADmobshyA D301A | This study |
|  | EP15 | EP3 pBADmobshyA H345A | This study |
|  | EP16 | EP3 pBADmobshyA H378A | This study |

**Supplementary Table 2.**
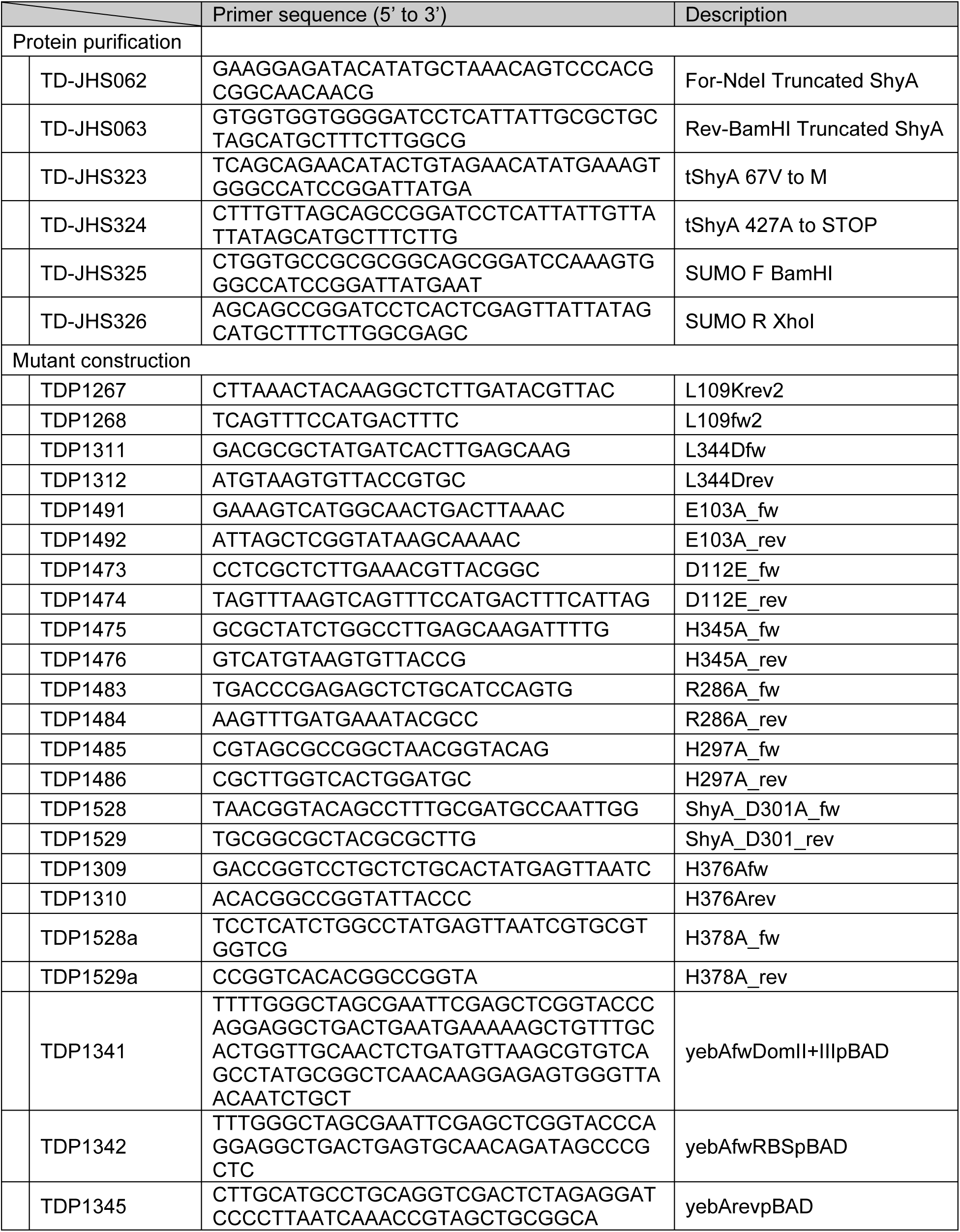
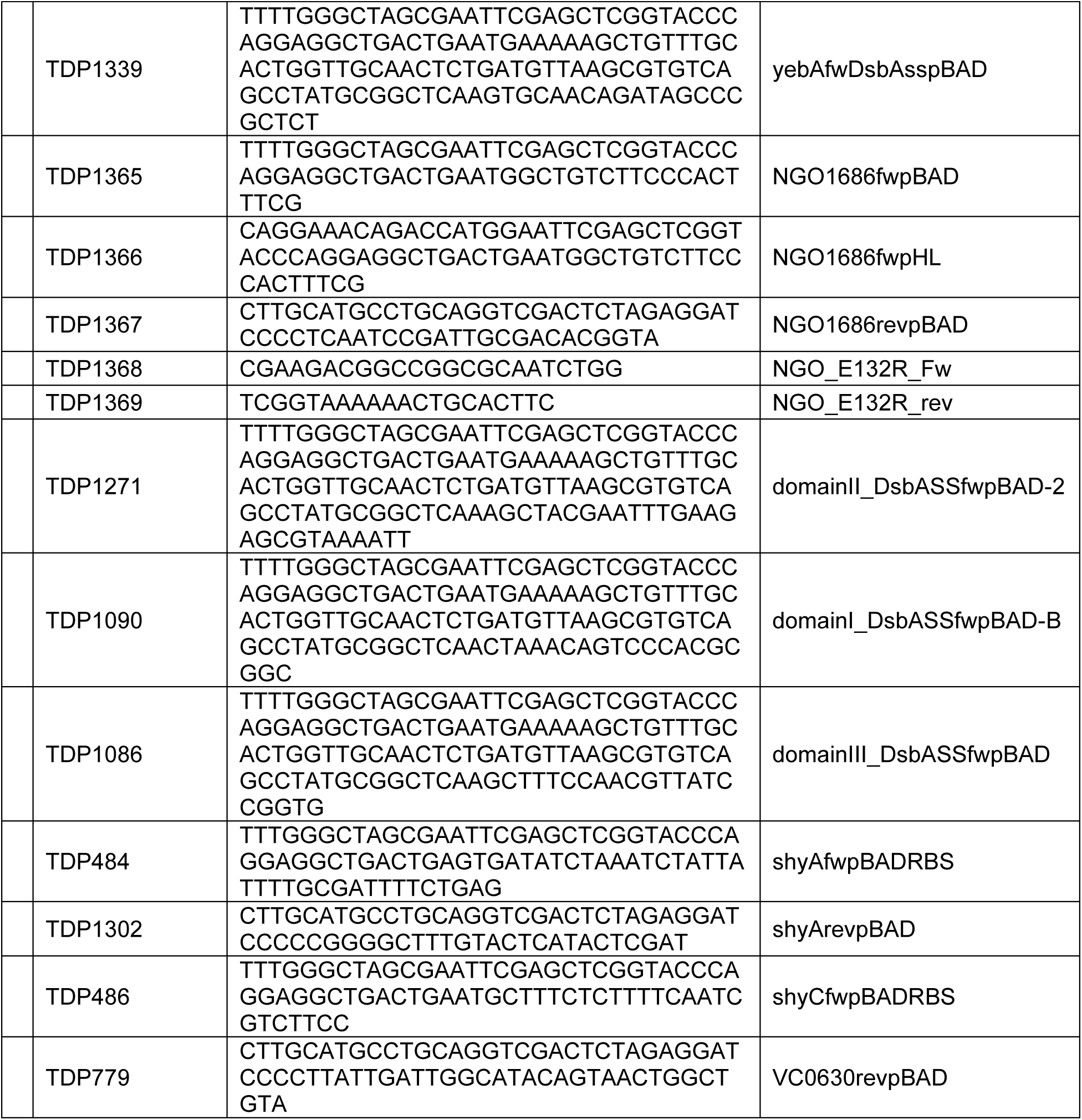
Oligonucleotides used in this study.

**Figure S1.**
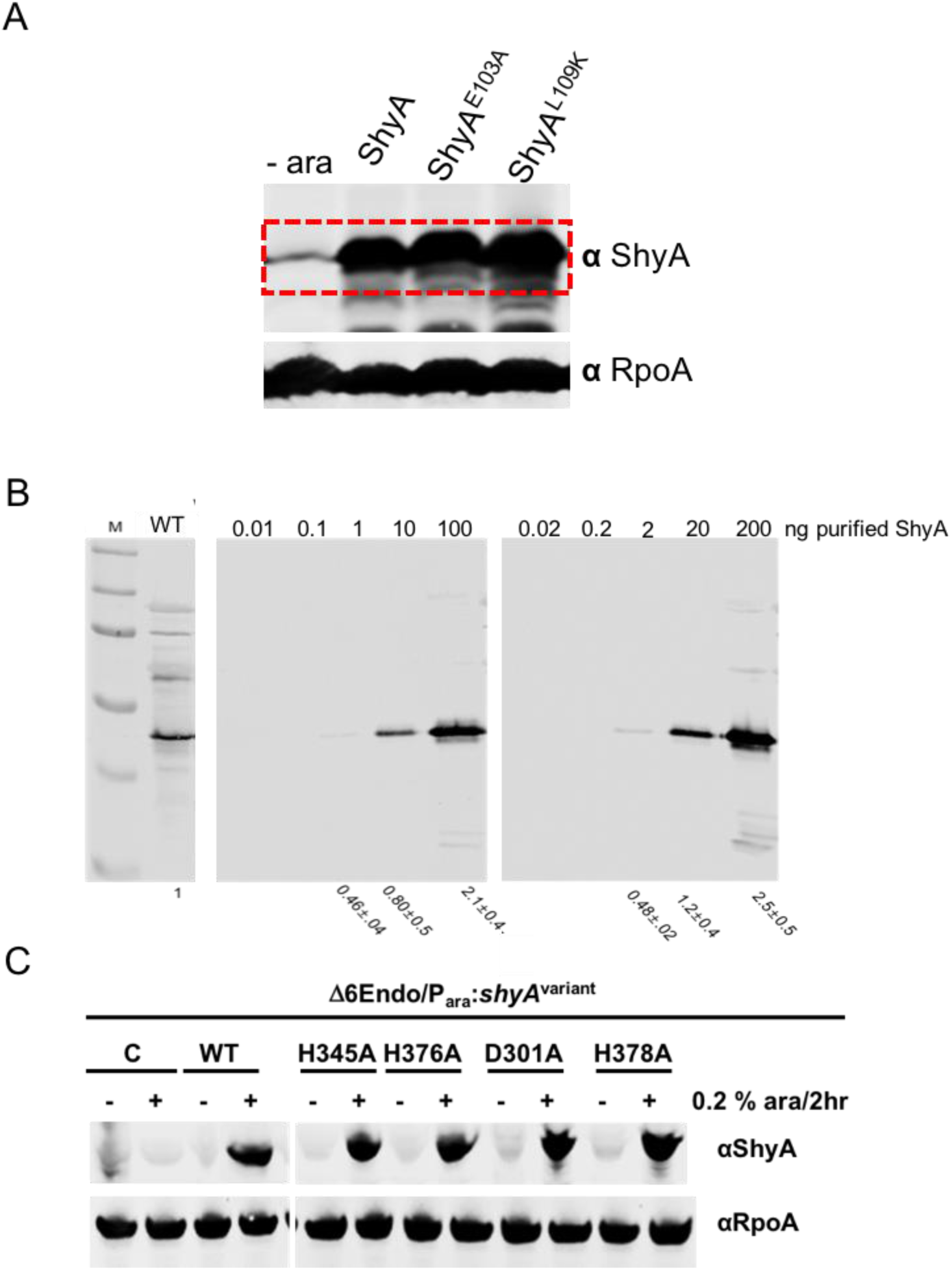
ShyA *in vivo* protein levels. **(A)** Cell lysates of Δ*shyA* strains carrying the respective ShyA variants under control of an arabinose-inducible promoter were subjected to Western Blot either before (–) or 2 hours after (+) addition of arabinose using polyclonal ShyA antibody. An RpoA antibody was used as loading control. **(B)** Estimation of native ShyA levels. Cell lysate (lane “WT”) and purified ShyA with known quantities (0.01 – 100 ng in the center gel, 0.02 – 200 ng in the right gel) were subjected to Western Blotting. The number of ShyA molecules/cell was estimated by comparing band intensities (numbers below gel image, normalized to wild-type levels) between known ShyA concentrations and the cell lysate obtained from a known number of cells. **(C)** Expression levels of catalytically inactive ShyA mutants. Cells were treated as described in **(A).** C = empty plasmid control.

**Figure S2.**
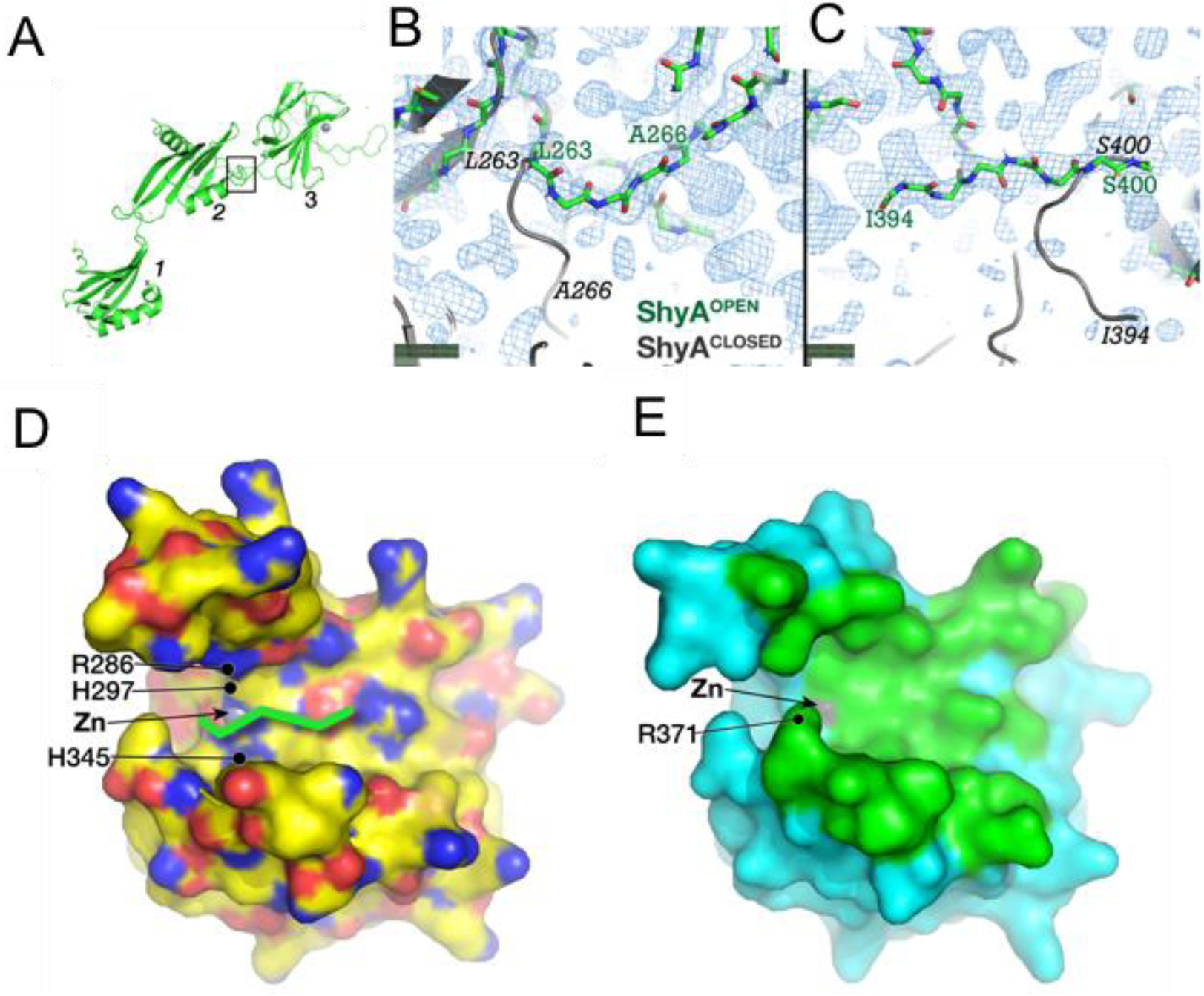
ShyA^OPEN^ interdomain linker polypeptides are clearly resolved in simulated annealing, composite omit electron density maps and take different paths compared to those in ShyA^CLOSED^. **(A)** Cartoon diagram of ShyA^OPEN^ (with domains labeled) with box outlining area of the omit maps depicted in **(B)** and **(C). (B)** Main-chain (green carbons) of ShyA^OPEN^ superposed on a composite omit difference electron density map. The dark grey cartoon (with italic labels) is the ShyA^CLOSED^ structure with Domain 2 superposed on Domain 2 of ShyA^OPEN^ showing how the folds diverge after L263. **(C)** The main-chain with green carbons represents the ShyA^OPEN^ polypeptide between I394 (end of Domain 3) and S400 (beginning of C-terminal helix). The path of the ShyA^CLOSED^ structure is shown in dark grey. **(D)** and **(E)** Putative substrate binding site on Domain 3 of ShyA. (**D**) Domain 3 of ShyA^OPEN^ with a solvent accessible surface colored by atom type: yellow, carbon; blue, nitrogen; red, oxygen. The zinc atom is a grey sphere. The green stick represents a hypothetical model of a peptidoglycan 4,3 crosslink: (from left-to-right) a D-Ala residue forming an amide bond to the N6 (sidechain Nε) of the *meso*-diaminopimelic acid. The position of the molecule is analogous to the substrate analog tetraglycine phosphinate co-crystallized with the M23 peptidase LytM (44). The scissile bond carbonyl (not shown) is pointed towards the Zn atom. (**E**) Solvent accessible surface of Domain 3 of ShyA^CLOSED^ structure. Green surface represents those atoms occluded by interactions with Domain 1 of the closed structure.

**Figure S3.**
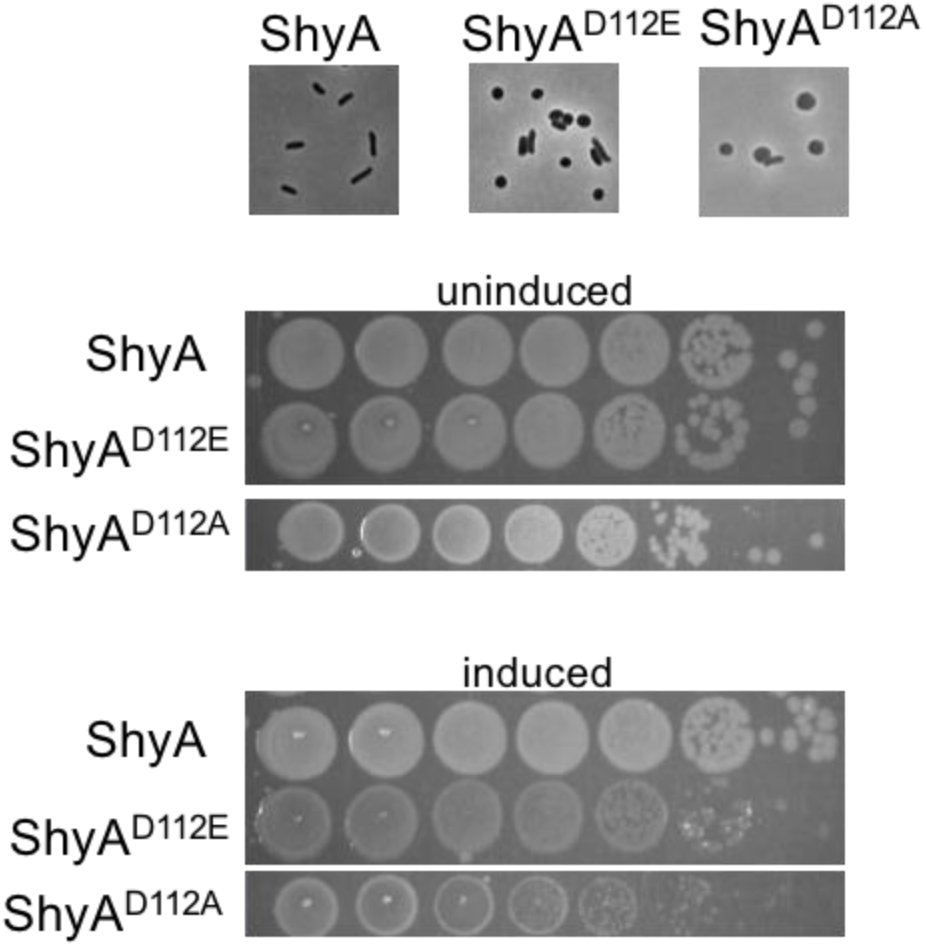
*In vivo* toxicity of ShyA^D112^ mutants. **(A)** ShyA or its D112E/A derivatives were overexpressed (0.2 % arabinose) in wild-type *V. cholerae* for 2 h, followed by visualization on an agarose pad**. (B)** Cells carrying arabinose-inducible ShyA or its D112E/A derivatives were plated on LB agar without (uninduced) or with (induced) 0.2 % arabinose.

**Fig. S4.**
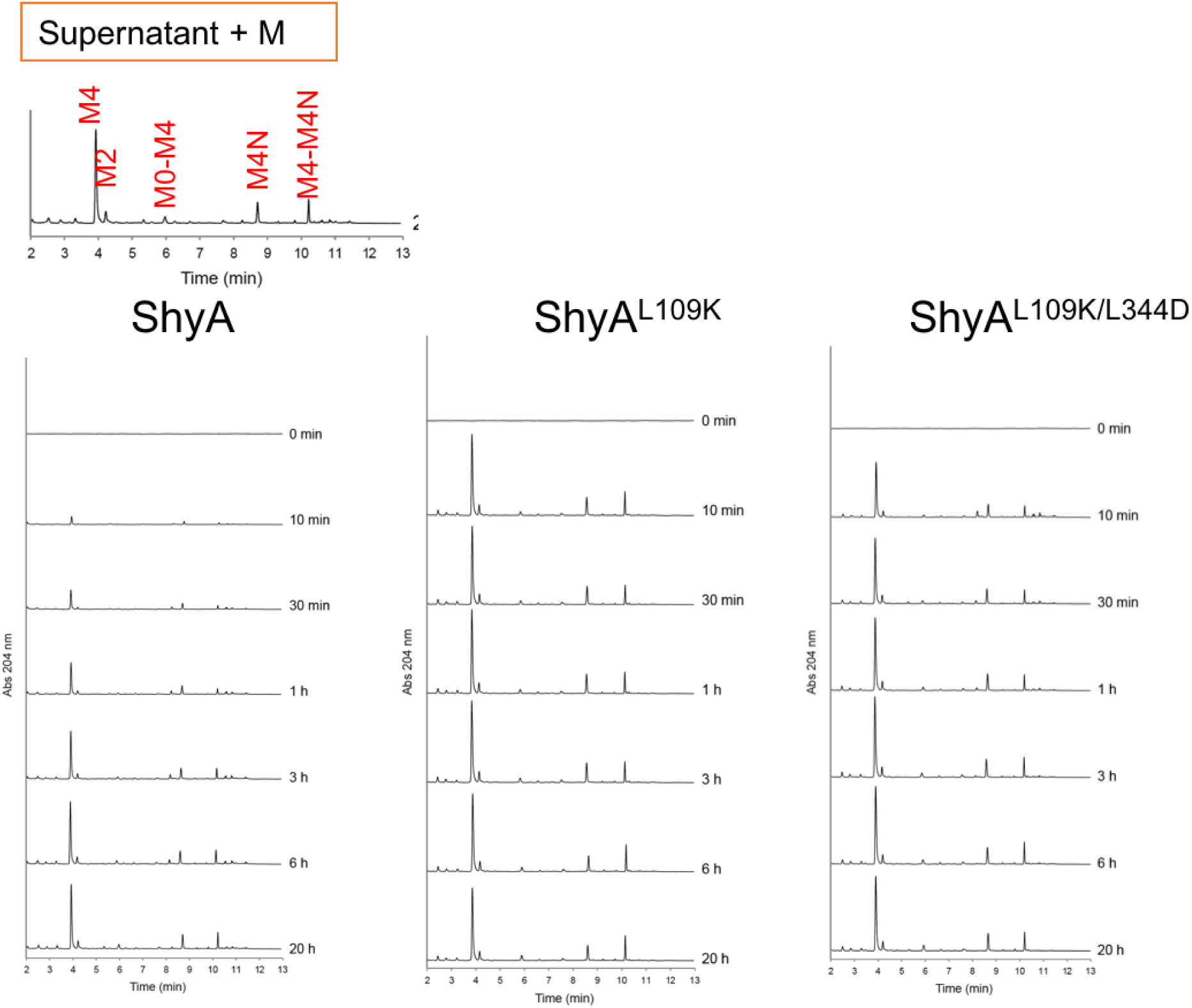
Chromatograms of ShyA activity experiments. Purified sacculi were digested as described in Methods. The supernatant of each reaction (*i.e.,* PG material solubilized by ShyA) was digested with muramidase (M) and then subjected to UPLC separation and detection by A_204_. Upper panel shows expected identity of peaks. M4, [*N*-acetylglucosamine (NAG)-*N*-acetylmuramic acid (NAM)]-tetrapeptide; M2, NAG-NAM-dipeptide; M0-M4, [NAG-NAM]-NAG-NAM-tetrapeptide; M4-M4N, [NAG-NAM]-tetrapeptide-[NAG-NAM]-tetrapeptide; N denotes termination in an anhydro-residue.

**Figure S5.**
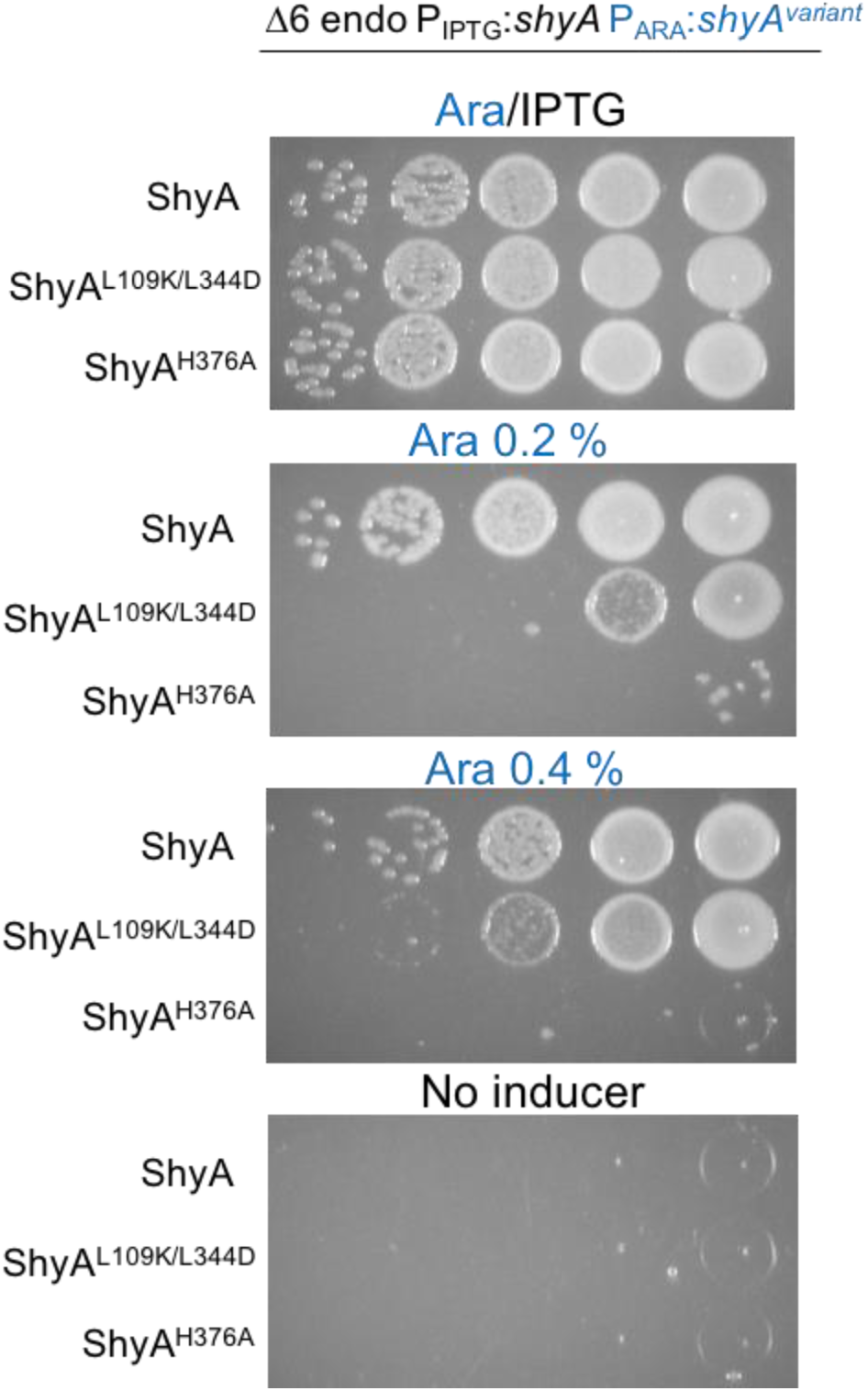
ShyA^L109K/L344D^ is partially functional. The Δ6 endo strain (Δ*shyABC* Δ*vc1537* Δ*tagE1/2* PIPTG:*shyA*) carrying plasmids expressing arabinose-inducible ShyA, its L109K/L344D or its H376A (inactive) derivatives were plated on growth medium containing 200 µM IPTG (inducing the chromosomal ShyA copy), 0.2 % arabinose (inducing plasmid-borne ShyA, ShyA^L109K/L344D^or ShyA^H376A^), 0.4% arabinose, or no inducer.

**Figure S6.**
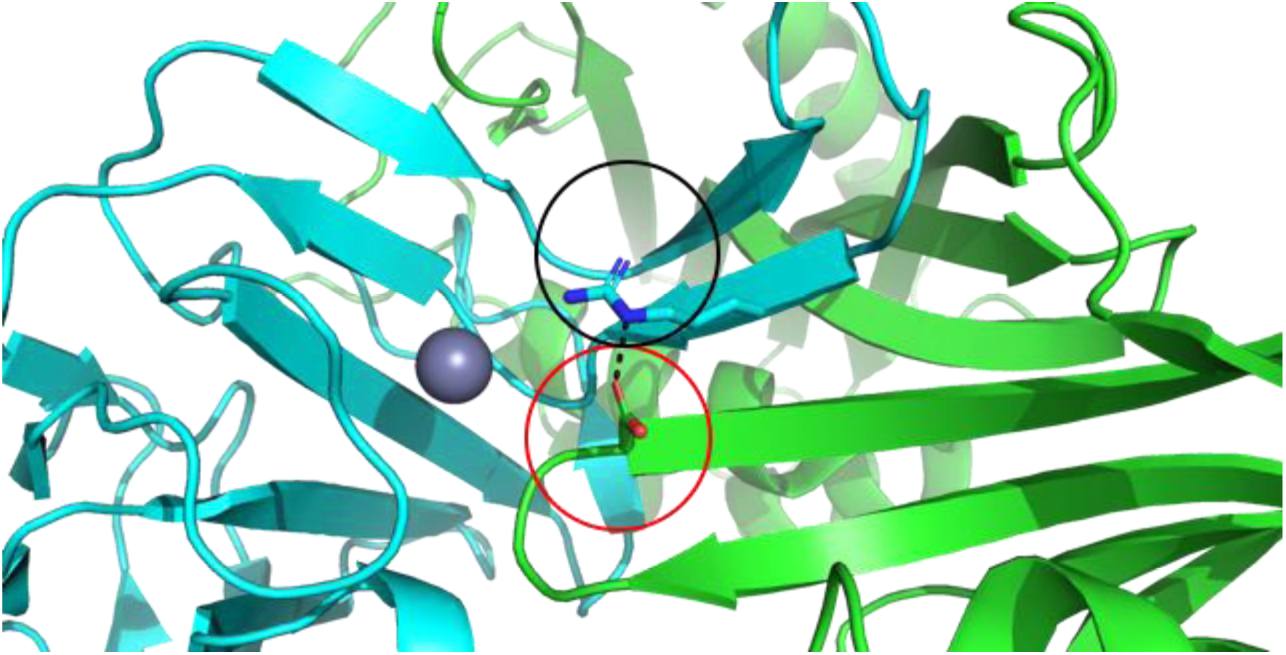
Residue E132 mediates putative Domain 1 – Domain 3 interactions in *Neisseria gonorrhea* NGO1686. Closeup view of the NGO1686 crystal structure [PDB 6MUK] with cartoon representation of putative Domain 1 (green) – Domain 3 (cyan) interface showing salt bridge between residues E132 (red circle) and R293 (black circle). Hydrogen bonding is represented with black dashed line and the grey sphere denotes the active site zinc.

**Figure S7.**
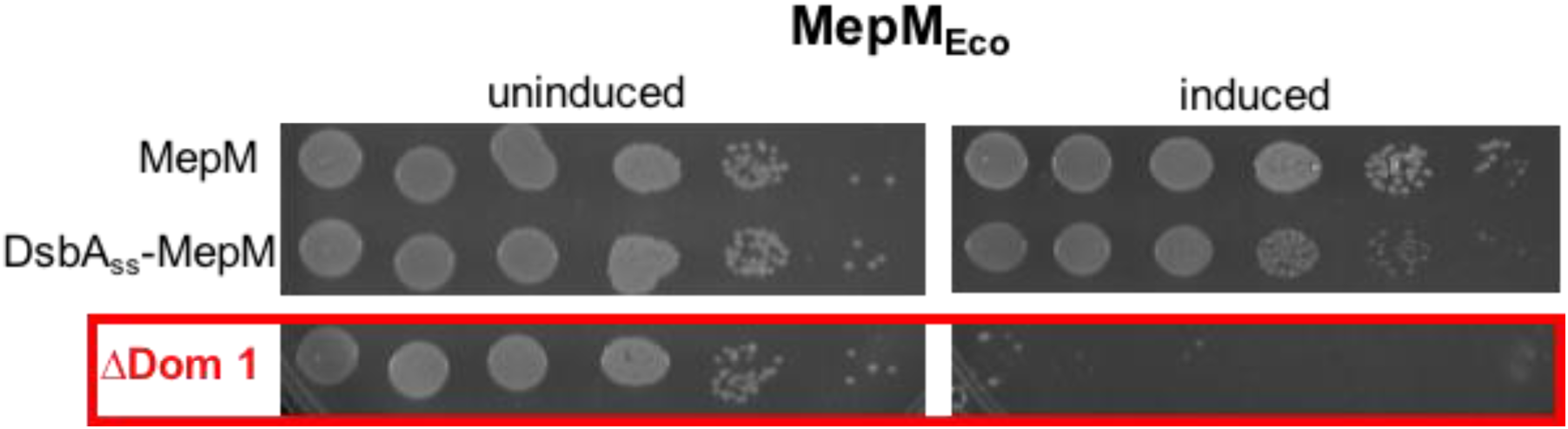
Overexpression toxicity of MepM^ΔDom1^. *E. coli* MG1655 carrying MepM or its DsbAss/ΔDomain 1 derivatives was diluted and spot-plated on LB agar containing either no inducer (“uninduced”) or 0.2 % arabinose (“induced”).

